# An N-degron proteolytic pathway modulates recipient susceptibility to T6SS DNase effectors

**DOI:** 10.64898/2026.04.17.719174

**Authors:** Yung-Hui Victoria Wen, Hsiao-Han Lin, Xing-Tai Zheng, Hau-Hsuan Hwang, Erh-Min Lai

## Abstract

The type VI secretion system (T6SS) is a contractile nanoweapon widely employed by Gram-negative bacteria to gain competitive advantages by injecting effector proteins into recipient cells. Although the biochemical activities of T6SS effectors have been well characterized, how recipient factors modulate effector toxicity remains poorly understood. Using *Agrobacterium* C58 as a model, previous work identified the *Escherichia coli* ClpAP protease as a recipient susceptibility (RS) factor that enhances T6SS-mediated interbacterial competition. *Agrobacterium* C58 deploys two DNase effectors, Tde1 and Tde2, as the major antibacterial weapon. Here, we demonstrate that the recipient ClpAP protease and its adaptor ClpS enhanced C58-mediated interbacterial competition in a Tde2-dependent manner in both intra- and interspecies competition. Ectopic expression of Tde2 in *E. coli* caused growth inhibition and DNA cleavage in the presence of a functional ClpAPS protease complex, but not in any of the *clpP, clpA* or *clpS* mutants. Notably, Tde2 accumulated in these mutants but not in wild-type cells, whereas a catalytic variant accumulated regardless of ClpAPS status, suggesting that Tde2 is not directly degraded by ClpAPS. Instead, Tde2 depends on ClpAPS for full toxicity, likely through degradation of inhibitory N-degron substrate(s). Affinity purification of His-tagged Tde2 in a *clpP* mutant background, followed by mass spectrometry, identified eight N-degron substrate candidates. Tde2-mediated interbacterial competition was significantly reduced by overexpression of three candidates. Among them, the Tde2 DNase domain directly associated with guanosine 5’-monophosphate reductase GuaC, supporting a model in which Tde2 toxicity is blocked by binding of GuaC. Collectively, our findings reveal an unanticipated layer of recipient-mediated regulation in T6SS competition and highlight proteolytic control of inhibitory substrates as a determinant of bacterial susceptibility during interbacterial conflict.

## Introduction

Competition for limited resources and space has driven bacteria to evolve diverse antagonistic strategies to survive in complex microbial communities (Ghoul and Mitri, 2016). Among these, Gram-negative bacteria frequently deploy the type VI secretion system (T6SS), a widespread contractile nanoweapon that enhances bacterial fitness by injecting toxic effector proteins into neighbouring recipient cells. By eliminating competitors, T6SS confers growth advantages in polymicrobial environments and contributes to niche establishment and persistence (Hood *et al*., 2010; Russell *et al*., 2014; Cianfanelli *et al*., 2016; Mijatović Scouten *et al*., 2025).

To withstand T6SS-mediated attack, recipient bacteria have evolved multiple defensive and counteroffensive strategies, including the production of exopolysaccharides that form protective physical barriers, the expression of cognate immunity proteins that neutralize incoming toxins, stress-responsive envelope remodeling, and retaliatory deployment of their own T6SS (Hernandez *et al*., 2020; Hersch *et al*., 2020).

In addition to these defense responses, recipient-derived components, termed recipient susceptibility (RS) factors, can paradoxically enhance intoxication and thereby increase vulnerability to T6SS attack (Lin *et al*., 2020a). In some cases, such factors promote toxin folding, activity, or entry into recipient cells. For example, the periplasmic disulfide bond oxidoreductase DsbA is required for the proper folding and activation of the *Serratia marcescens* peptidoglycan hydrolase effectors Ssp2 and Ssp4 after delivery into recipient cells (Mariano *et al*., 2018). Recipient surface structures can likewise act either as protective barriers or as RS determinants. A CRISPR interference (CRISPRi)-based screen identified MsbA, a component of lipopolysaccharide (LPS) biogenesis in *Pseudomonas aeruginosa*, as a recipient determinant required for the cytotoxic activity of the T6SS amidase Tae1 (Trotta *et al*., 2023), whereas in *Salmonella enterica* LPS acts as a barrier to T6SS-mediated attack (Tsai *et al*., 2025). Comparable recipient dependencies have also been described for contact-dependent inhibition (CDI) systems, in which the *Escherichia coli* type Vb secretion toxin CdiA-CT^EC93^ engages specific outer membrane receptors, such as BamA and AcrB, to enable toxin entry into recipient cells (Aoki *et al*., 2005; Aoki *et al*., 2008; Mariano *et al*., 2018).

In soil-dwelling phytopathogen agrobacteria, the T6SS can function as an antibacterial weapon that promotes interbacterial competition during plant colonization (Ma *et al*., 2014; Wu *et al*., 2019) and contributes to disease-associated fitness (Wang *et al*., 2023). *Agrobacterium* strain C58 encodes three effectors, in which two DNase effectors (Tde1 and Tde2) serve as the primary antibacterial effectors. Its interbacterial activity extends across both intra- and interspecies contexts, targeting diverse bacteria including *Agrobacterium* species, *E. coli*, *P. aeruginosa*, *Dickeya dadantii* and *Sphingomonas* spp. (Ma *et al*., 2014; Bondage et al., 2016; Wu *et al*., 2019; Wu *et al*., 2020; Santos *et al*., 2020; Wang *et al*., 2023; Santos *et al*., 2024). Using the *E. coli* BW25113 Keio knockout collection as recipients in competition assays with *Agrobacterium* C58, our previous work identified eighteen mutants with increased resistance to T6SS-mediated interbacterial competition (Lin *et al*., 2020b). Among these, the unfoldase ClpA, but not ClpX, was required together with the protease ClpP to enhance recipient susceptibility to *Agrobacterium* C58 T6SS-mediated interbacterial competition. ClpAP is a conserved proteolytic complex involved in protein quality control (Katayama-Fujimura *et al*., 1987; Katayama-Fujimura *et al*., 1990; Wong and Houry, 2004; Olivares *et al*., 2015) and degrades ssrA-tagged proteins (Gottesman *et al*., 1998) as well as N-end rule (N-degron) substrates via the ClpS adaptor (Dougan *et al*., 2002; Zeth *et al*., 2002). These findings raise the possibility that intrinsic recipient proteostasis pathways can shape the outcome of T6SS-mediated interbacterial competition.

Here, we reveal that *Agrobacterium* harnesses the recipient ClpAPS protease system to degrade Tde2-associated inhibitors, thereby mitigating their antagonistic effects and facilitating Tde2-induced DNA degradation. We further identify eight Tde2-associated N-degron substrates, one of which, guanosine 5’-monophosphate reductase (GuaC), associates with the Tde2 DNase domain and attenuates its toxicity. Together, these findings support a model in which a recipient proteolytic pathway promotes susceptibility to a T6SS effector by degrading effector-associated inhibitory factors.

## Results

### *Agrobacterium* Tde2-mediated interbacterial competition depends on the recipient ClpAPS protease complex

Previous work showed that C58 T6SS-mediated interbacterial competition depends on the *E. coli* recipient susceptibility (RS) factors ClpA and ClpP (Lin *et al*. 2020b). We therefore asked whether this ClpAP-dependent susceptibility is linked to specific C58 T6SS effectors. In *Agrobacterium* C58, Tde1 and Tde2 belong to the Ntox15 superfamily and share a conserved HxxD catalytic motif required for DNase activity, and function as the major antibacterial effectors (Ma *et al*., 2014). We performed interbacterial competition assays using C58 WT and derivative attacker strains delivering Tde1 (Δ*tdei2*), Tde2 (Δ*tdei1*), or neither effector (Δ*tdei*), as well as a strain expressing a DNase-deficient Tde2 variant carrying H439A and D442A substitutions, Δ*tdei*(*tde2^HD^*-*tdi2*). These strains were tested against *E. coli* BW25113 wild-type (BW WT) and the indicated *clp* mutants (Δ*clpP*, Δ*clpA*, Δ*clpS* and Δ*clpX*) as recipients. The Δ*clpS* was included to assess the involvement of the ClpAP adaptor ClpS in this susceptibility, whereas the Δ*clpX* served as a negative control. T6SS-mediated interbacterial competition was quantified using the susceptibility index (SI), defined as the log_10_ difference in colony-forming units (CFU) recovered from recipient bacteria co-cultured with the Δ*tdei* negative-control attacker versus the indicated C58 strains, with higher SI values indicating greater susceptibility (Lin *et al*. 2020b).

C58 WT, Δ*tdei2* and Δ*tdei1* showed similar killing of the BW WT recipient while no killing activity was observed by Δ*tdei* (Fig. 1A, S1), indicating delivery of either Tde1 or Tde2 is sufficient for full T6SS killing activity. By contrast, C58 WT and Δ*tdei2* showed reduced killing against the Δ*clpP*, Δ*clpA* and Δ*clpS* recipients, but not the Δ*clpX* recipient, relative to BW WT, although only Δ*clpP* showed a statistically significant difference. Strikingly, Δ*tdei1*, the Tde2-mediated attacker, was almost completely impaired in killing the Δ*clpP*, Δ*clpA* and Δ*clpS* recipients. No killing was observed for the Δ*tdei*(*tde2^HD^*-*tdi2*) attacker against any recipient strain (Fig. 1A). These findings indicate that the contribution of recipient ClpAPS to C58 T6SS-mediated interbacterial competition is primarily linked to Tde2.

**Figure 1.**
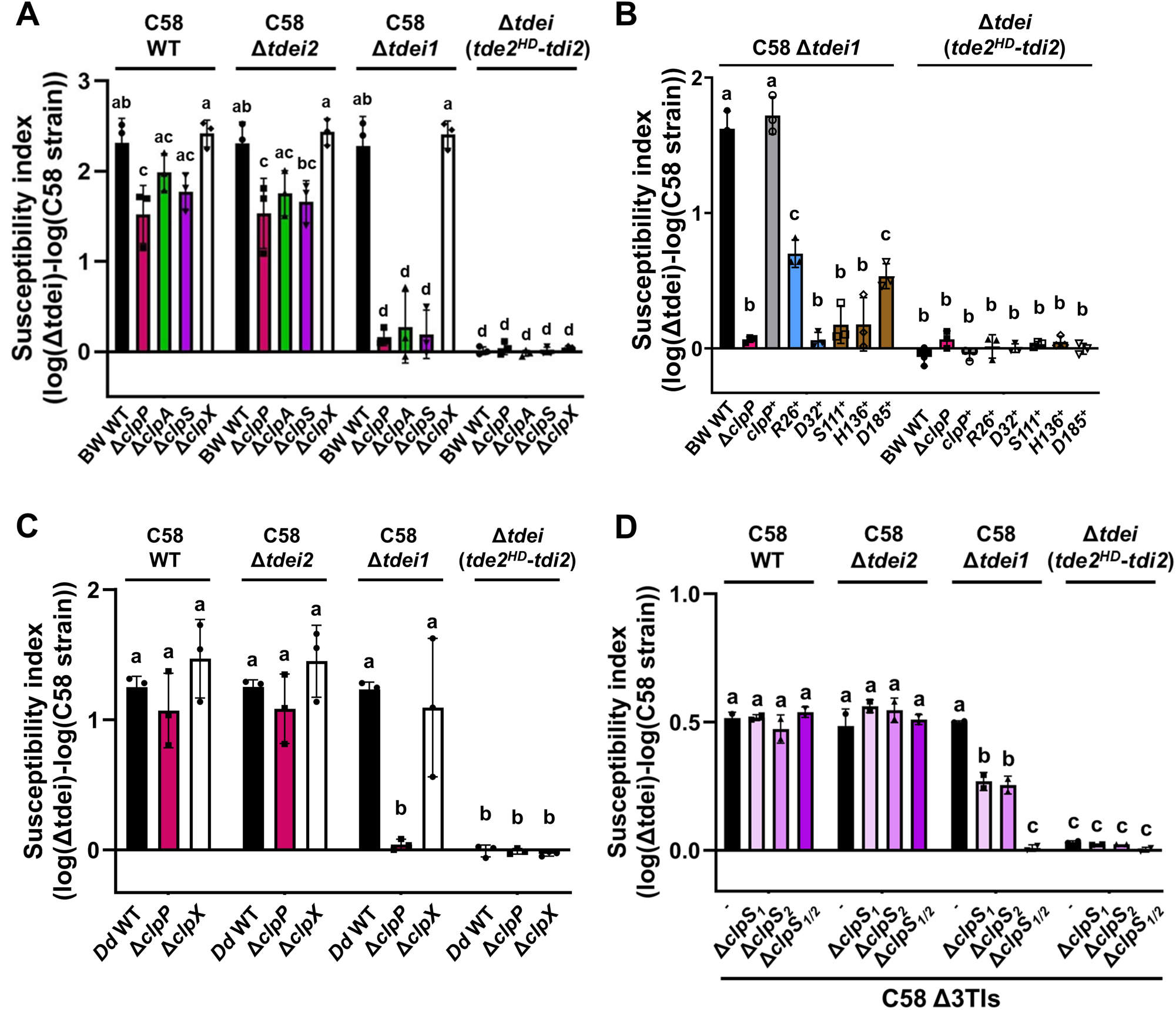
*Agrobacterium* type VI secretion system (T6SS) Tde2-mediated interbacterial competition depends on the recipient ClpAPS complex. *Agrobacterium* C58 attacker strains indicated above each plot were co-cultured with the indicated recipient strains (bottom) at a 10:1 ratio for 16 h at 25 °C. T6SS-mediated interbacterial competition effect was quantified as the susceptibility index (SI), defined as the log_10_ of the CFU count from the recipient cells co-cultured with C58 Δ*tdei* minus the log_10_ of the CFU count from the recipient cells co-cultured with other C58 strains. The data are mean ± SD from three biological replicates. Statistical significance was determined by two-way analysis of variance (ANOVA) followed by Tukey’s honestly significant difference (HSD) test (*p* < 0.05); groups with different letters differ significantly. **A.** Susceptibility of *Escherichia coli* BW25113 WT (BW WT), Δ*clpP*::Kan^R^ (Δ*clpP*), Δ*clpA*::Kan^R^ (Δ*clpA*), Δ*clpS*::Kan^R^ (Δ*clpS*) and Δ*clpX*::Kan^R^ (Δ*clpX*), each carrying pTrc200-HA. **B.** Susceptibility of *E. coli* BW25113 WT carrying pTrc200-HA (BW WT), Δ*clpP* carrying pTrc200-HA or its derivatives expressing WT ClpP (*clpP*^+^) and its variants ClpP^R26A^ (R26^+^), ClpP^D32A^ (D32^+^), ClpP^S111A^ (S111^+^), ClpP^H136A^ (H136^+^) or ClpP^D185A^ (D185*^+^*). **C.** Susceptibility of *Dickeya dadantii* strain 3937 WT (*Dd* WT), Δ*clpP*::Kan^R^ (Δ*clpP*) and Δ*clpX*::Kan^R^ (Δ*clpX*), each carrying pTrc200-HA. **D.** Susceptibility of *Agrobacterium* C58 Δ3TIs (null-effector mutant), Δ3TIsΔ*clpS_1_* (Δ*clpS_1_*), Δ3TIsΔ*clpS_2_* (Δ*clpS_2_*) and Δ3TIsΔ*clpS_1/2_* (Δ*clpS_1/2_*), each carrying pTrc200-HA.

Lin *et al*. (2020b) showed that C58 T6SS-mediated interbacterial competition depends on both ClpA-ClpP complex formation and ClpP protease activity. We therefore investigated whether Tde2-mediated interbacterial competition similarly requires a functional ClpAP protease complex. To test this, we used *E. coli* recipient strains expressing WT ClpP or ClpP variants defective in ClpA binding (R26A^+^ and D32A^+^) (Bewley *et al*., 2006) or protease activity (S111A^+^, H136A^+^ and D185A^+^) (Maurizi *et al*., 1990; Wang *et al*., 1997) for interbacterial competition assays. As expected, expression of WT ClpP in the Δ*clpP* fully restored susceptibility to the BW WT level (Fig. 1B). By contrast, the R26A variant, which retains partial ClpA-binding activity, conferred only partial restoration, whereas the D32A variant, which is defective in ClpA binding, remained as resistant as the Δ*clpP* (Fig. 1B). Likewise, all three catalytic triad mutants conferred greater resistance than WT ClpP, and the D185A variant, which retains weak protease activity *in vitro* (Lin *et al*., 2020b), showed partial restoration (Fig. 1B). Together, these results suggest that Tde2-mediated interbacterial competition depends on both ClpA-ClpP complex formation and ClpP protease activity in recipient cells.

Whereas expression of ClpP or ClpS restored susceptibility to the Δ*tdei1* attacker in the corresponding deletion mutant recipients (*clpP^+^* and *clpS^+^*) (Fig. 1B, S2A), expression of ClpA in the Δ*clpA* recipient (*clpA^+^*) failed to restore susceptibility to Tde2-mediated interbacterial competition (Fig. S2B). Because the complementation constructs were expressed from the IPTG-inducible *trc* promoter on pTrc200-HA, and both ClpS and ClpA accumulated to levels higher than those of the corresponding endogenous proteins (Fig. S2C, S2D), the failure of ClpA complementation was unlikely to reflect insufficient ClpA expression. Instead, we explain that excess ClpA might exert a dominant-negative effect. Omitting IPTG induction still failed to restore susceptibility in the Δ*clpA*, likely because ClpA levels remained more abundant than the endogenous level (Fig. S2B, S2D).

We next assessed whether the requirement for recipient ClpAPS in Tde2-mediated interbacterial competition is conserved. To address this, we first tested *Dickeya dadantii* strain 3937 (Li *et al*., 2010), a γ-proteobacterial plant pathogen that causes soft-rot disease and belongs to the same class as *E. coli*, as the recipient. Tde2-mediated interbacterial competition was significantly reduced against the *D. dadantii* Δ*clpP* compared with WT, whereas the Δ*clpX* remained as susceptible as WT (Fig. 1C). We then extended this analysis to an α-proteobacterial recipient using the susceptible *Agrobacterium* C58 Δ3TIs strain, which lacks all three toxin-immunity pairs (Ma *et al*., 2014). The C58 genome encodes three *clpP* genes (*clpP_1_*, *clpP_2_* and *clpP_3_*), one *clpA* and two *clpS* genes (*clpS_1_* and *clpS_2_*). We successfully generated single- and double-deletion *clpP* mutants, but were unable to obtain a triple *clpP* mutant. We also constructed *clpS* single and double mutants in the C58 Δ3TIs background. While no clear increase in resistance was observed in C58 Δ3TIs derivatives carrying additional single or double *clpP* deletions during C58 T6SS-mediated interbacterial competition (Fig. S3), Tde2-mediated interbacterial competition was reduced in the *clpS* mutants, with Δ*clpS_1/2_* showing significantly greater resistance than the Δ3TIs control (Fig. 1D). Collectively, these findings suggest that Tde2 depends on recipient ClpAPS during interbacterial competition across both intra- and interspecies interactions.

### Tde2 DNase-induced toxicity requires ClpAPS for growth inhibition and DNA cleavage in recipient cells

To investigate whether Tde2-mediated DNase toxicity depends directly on the recipient ClpAPS, we ectopically expressed His-tagged WT Tde2 and its DNase-deficient mutant (Tde2^HD^) from the L-arabinose-inducible promoter on pJN105 (Newman and Fuqua, 1999). These constructs were introduced into *E. coli* BW25113 WT and the indicated *clp* mutants, and their effects were assessed by growth analysis and DNA degradation assays. As anticipated, induction of Tde2 expression inhibited growth in WT and Δ*clpX*, as compared to the empty vector (EV) control. However, Tde2-induced growth inhibition was fully rescued in the Δ*clpP* and partially relieved in the Δ*clpA* and Δ*clpS*, but not in the Δ*clpX* (Fig. 2A). Expression of Tde2^HD^ had no detectable effect on growth in any strain (Fig. 2A). These results suggest that Tde2 depends on the recipient ClpAPS to exert DNase-induced growth inhibition.

**Figure 2.**
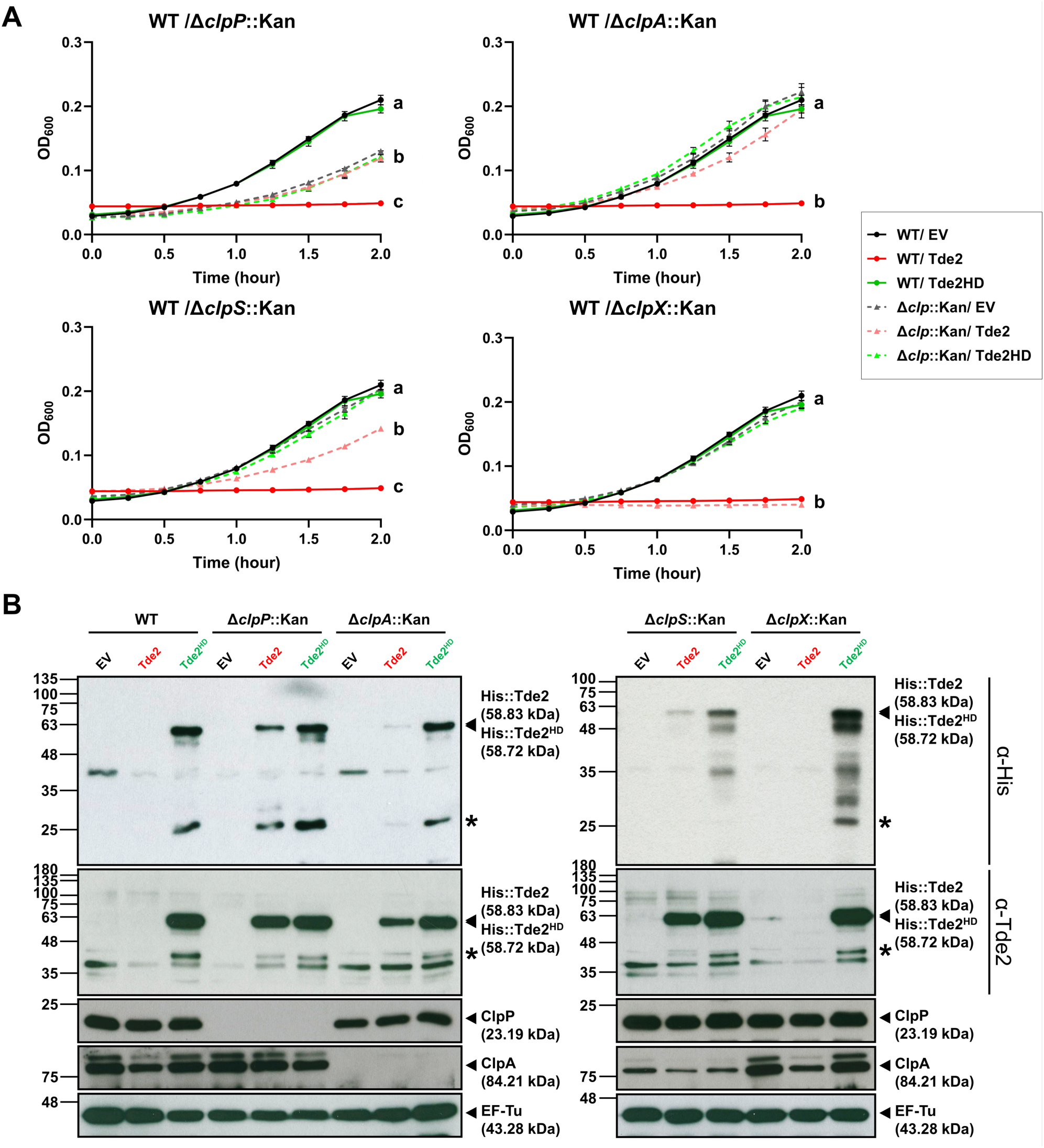
Tde2 DNase-induced toxicity depends on the recipient ClpAPS complex. **A.** Growth of *Escherichia coli* BW25113 WT (solid line) and the indicated *clp* mutants (dashed line) carrying empty vector (EV, black) or plasmid expressing His-tagged Tde2 (red) or catalytically inactive Tde2HD (green). Cultures were supplemented with 0.005% L-arabinose at 0 h. The data represent mean ± SD from six biological replicates. Statistical significance was determined by one-way analysis of variance (ANOVA) followed by Tukey’s honestly significant difference (HSD) test (*p* < 0.05); groups with different letters differ significantly. **B.** Immunoblot analysis of samples from the growth inhibition assay in **A**, probed with antibodies against His, Tde2, ClpP, ClpA, and EF-Tu. Molecular weight markers are shown on the left, and the predicted molecular masses of the indicated proteins are shown on the right. An asterisk (*) denotes truncated forms of Tde2 or Tde2^HD^.

We next determined whether the loss of Tde2 DNase-induced toxicity in the Δ*clpP*, Δ*clpA* and Δ*clpS* was associated with altered protein abundance or reduced DNase activity. By examining Tde2 protein levels in BW WT and the *clp* mutants, unexpectedly, Tde2 accumulated to much higher levels in the Δ*clpP*, Δ*clpA* and Δ*clpS* than in WT and Δ*clpX*, in which it was barely detectable. Tde2^HD^ accumulated to even higher levels and also resulted in protein truncations (Fig. 2B). Thus, the reduced Tde2 DNase-induced toxicity in the absence of an intact ClpAPS cannot be explained by Tde2 protein level.

We next examined whether Tde2 DNase-induced toxicity correlated with DNA degradation. Plasmid DNA degradation was detected in WT and the Δ*clpX* expressing Tde2, but not in the Δ*clpP*, Δ*clpA* and Δ*clpS* (Fig. 3A). This effect depended on Tde2 DNase activity, as no DNA degradation was observed in any strain expressing Tde2^HD^ (Fig. 3A). To test more directly whether Tde2 induces DNA breaks, we performed TUNEL assays and quantified TUNEL-positive cells by flow cytometry following FITC-dUTP labeling of 3’-hydroxyl DNA ends. TUNEL-positive populations were defined relative to the unstained control for each strain. Tde2 expression yielded TUNEL-positive signals in 50-60% of WT cells, but in only 1-7% of Δ*clpP* cells and 5-11% of Δ*clpS* cells, while little or no signal was detected in strains carrying EV or expressing Tde2^HD^ (Fig. 3B).

**Figure 3.**
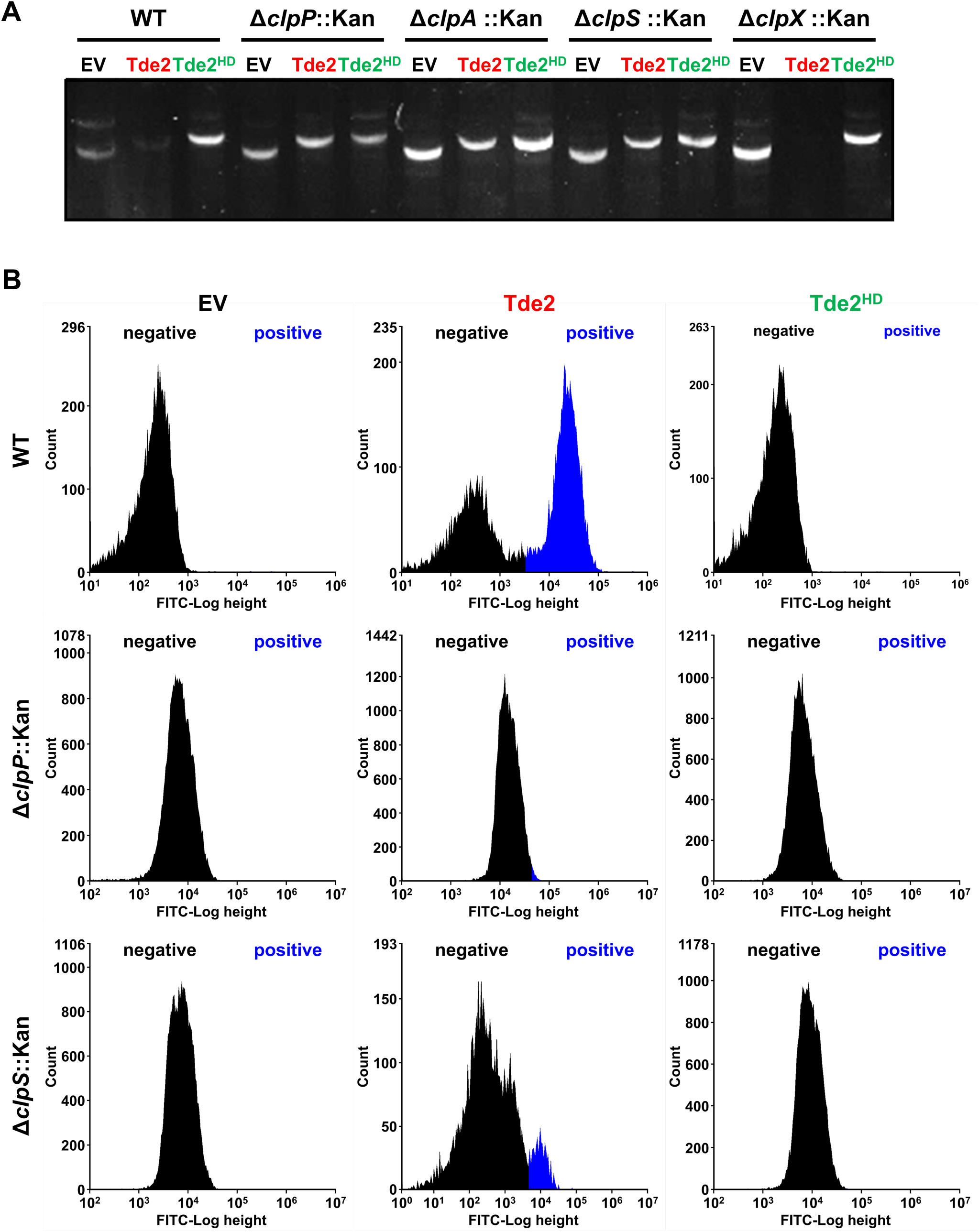
Tde2 DNase-induced DNA damage depends on the recipient ClpAPS complex. *Escherichia coli* BW25113 WT and the indicated *clp* mutants carrying empty vector (EV) or plasmid expressing His-tagged Tde2 or catalytically inactive variant Tde2^HD^ were induced with 0.005% L-arabinose and analyzed for DNA damage. **A.** Agarose gel electrophoresis of plasmid DNA extracted from the growth inhibition assay sample. **B.** TUNEL assay of cells collected from the same cultures and analyzed by flow cytometry. FITC fluorescence intensity is shown on the x-axis and cell count on the y-axis. Black indicates TUNEL-negative populations and blue indicates TUNEL-positive populations.

Together, these findings demonstrated that Tde2 DNase-induced toxicity and DNA cleavage require the ClpAPS. Although Tde2 accumulated in the absence of ClpAPS, its DNase-associated toxicity was markedly reduced, indicating that the ectopically expressed Tde2 to exert its cytotoxic DNase activity depends on an intact ClpAPS system.

### Identification of N-degron substrates associated with Tde2

Given that Tde2 DNase-mediated toxicity requires a functional ClpAPS protease complex and that inversely correlates with Tde2 protein abundance, we asked whether Tde2 itself is a direct substrate of ClpAPS to activate its toxicity. However, ClpAPS functions as a processive protease rather than through a limited proteolytic reaction, and no truncated Tde2 were detected in BW WT. A truncated Tde2^HD^ could be detected in both WT and *clp* mutants, suggesting that Tde2 is unlikely to be the relevant ClpAPS substrate. Thus, we hypothesized that *Agrobacterium* C58 exploits the recipient ClpAPS protease system to degrade the inhibitory factors that associate with Tde2 and suppress its DNase activity. Because ClpAPS protease complex degrades N-end rule (N-degron) substrates in bacteria (Dougan *et al*., 2002; Zeth *et al*., 2002), these inhibitory factors could be the N-degron substrates.

If this hypothesis is correct, Tde2 should form a complex with such substrates in the absence of ClpP, ClpA or ClpS. To test this, we expressed His-tagged Tde2 in BW25113 Δ*clpP*, isolated His-Tde2 and associated proteins by Ni-NTA pull-down, and identified Tde2-associated proteins by liquid chromatography-tandem mass spectrometry (LC-MS/MS). Empty vector and His-tagged sfGFP were included as negative controls. Across five biological replicates of two independent experiments, eight proteins that have been reported or proposed to be N-degron substrates (Humbard *et al*., 2013) were enriched in the His-Tde2 sample relative to both negative controls (Table 1). Thus, they were selected for further investigation.

**Table 1.**
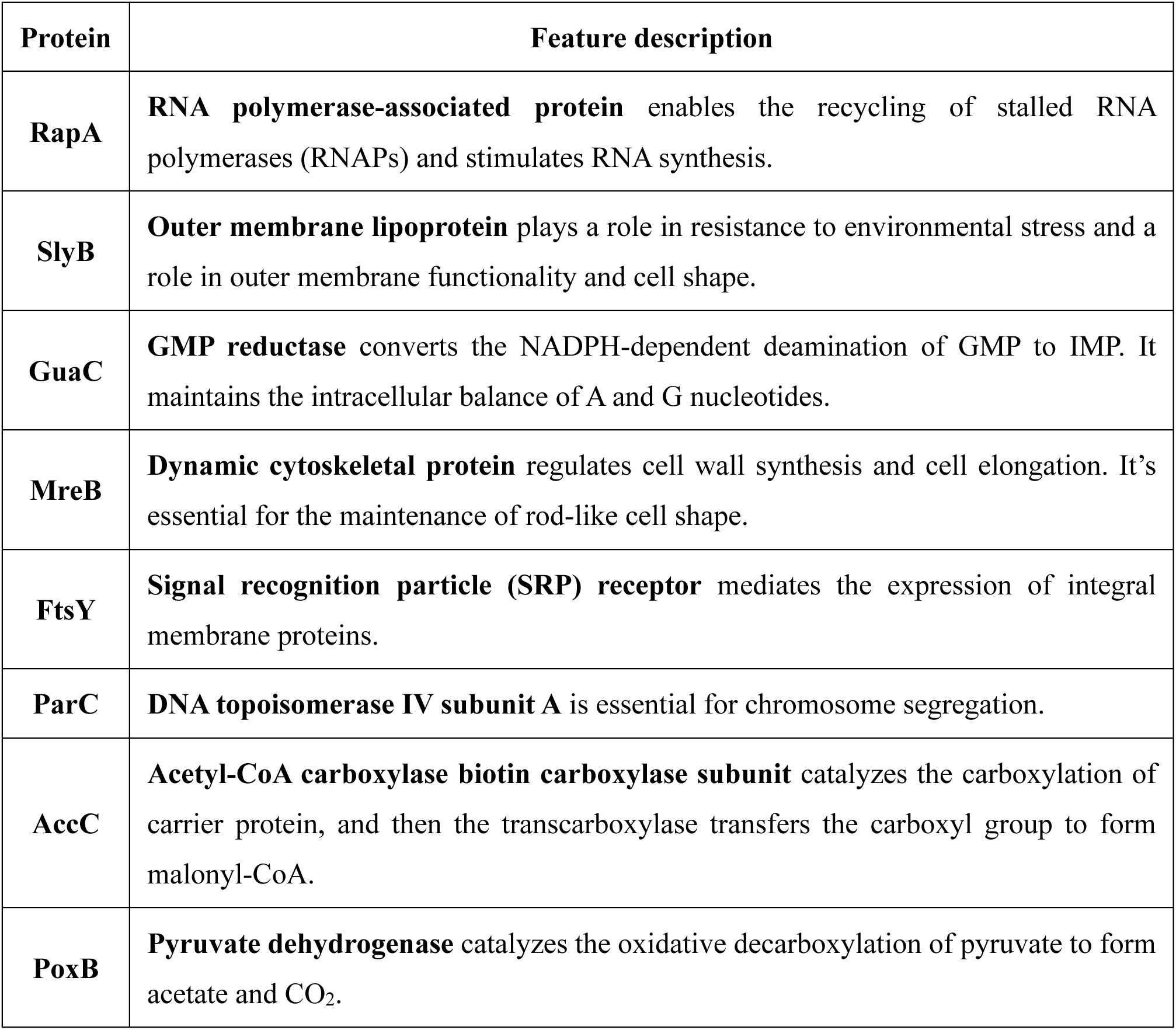
LC–MS/MS analysis identifies candidate N-degron substrates among Tde2-associated proteins in the *clpP* mutant background.

### Tde2-mediated interbacterial competition is reduced by overexpression of GuaC, AccC, or PoxB in recipient cells

We reasoned that the eight Tde2-associated N-degron substrates (RapA, SlyB, GuaC, MreB, FtsY, ParC, AccC and PoxB) may bind C58-delivered Tde2 and inhibit its DNase toxicity. To test whether these candidates modulate Tde2-mediated interbacterial competition, we performed interbacterial competition assays using the respective *E. coli* deletion mutants or overexpression strains. Several candidate genes are essential and therefore could not be deleted. In addition, overexpression of *rapA* or *mreB* impaired growth on minimal AK agar, the medium used for interbacterial competition assays, and these strains were therefore excluded from further analysis. We therefore examined the effects of the remaining *E. coli* deletion mutants (Δ*rapA*, Δ*slyB*, Δ*guaC* and Δ*poxB*) and overexpression strains (WT+*slyB*, WT+*guaC*, WT+*ftsY*, WT+*parC*, WT+*accC* and WT+*poxB*) on C58 T6SS-mediated killing.

Competition assays revealed that both C58 WT and the Δ*tdei2* attackers caused similar killing of the Δ*rapA*, Δ*slyB* and Δ*guaC* recipients relative to BW WT, whereas the Δ*tdei1* attacker showed reduced killing only of the Δ*poxB* recipient (Fig. 4). The absence of phenotypes in several deletion mutants may reflect functional redundancy among multiple Tde2-associated inhibitory factors. Thus, the overexpression approach is a feasible approach to test their effects. We show that the Δ*tdei1* attacker exhibited reduced killing of recipients overexpressing GuaC or PoxB relative to BW WT, whereas AccC overexpression reduced susceptibility to either the Tde1 or Tde2-mediated attackers (Fig. 5). By contrast, overexpression of SlyB, FtsY or ParC did not alter killing by C58 T6SS effectors (Fig. 5). Together, these findings suggest that recipient GuaC specifically impairs Tde2-mediated interbacterial competition, whereas AccC may attenuate both Tde1 and Tde2. PoxB, however, may additionally influence C58 T6SS-mediated interbacterial competition through different mechanisms.

**Figure 4.**
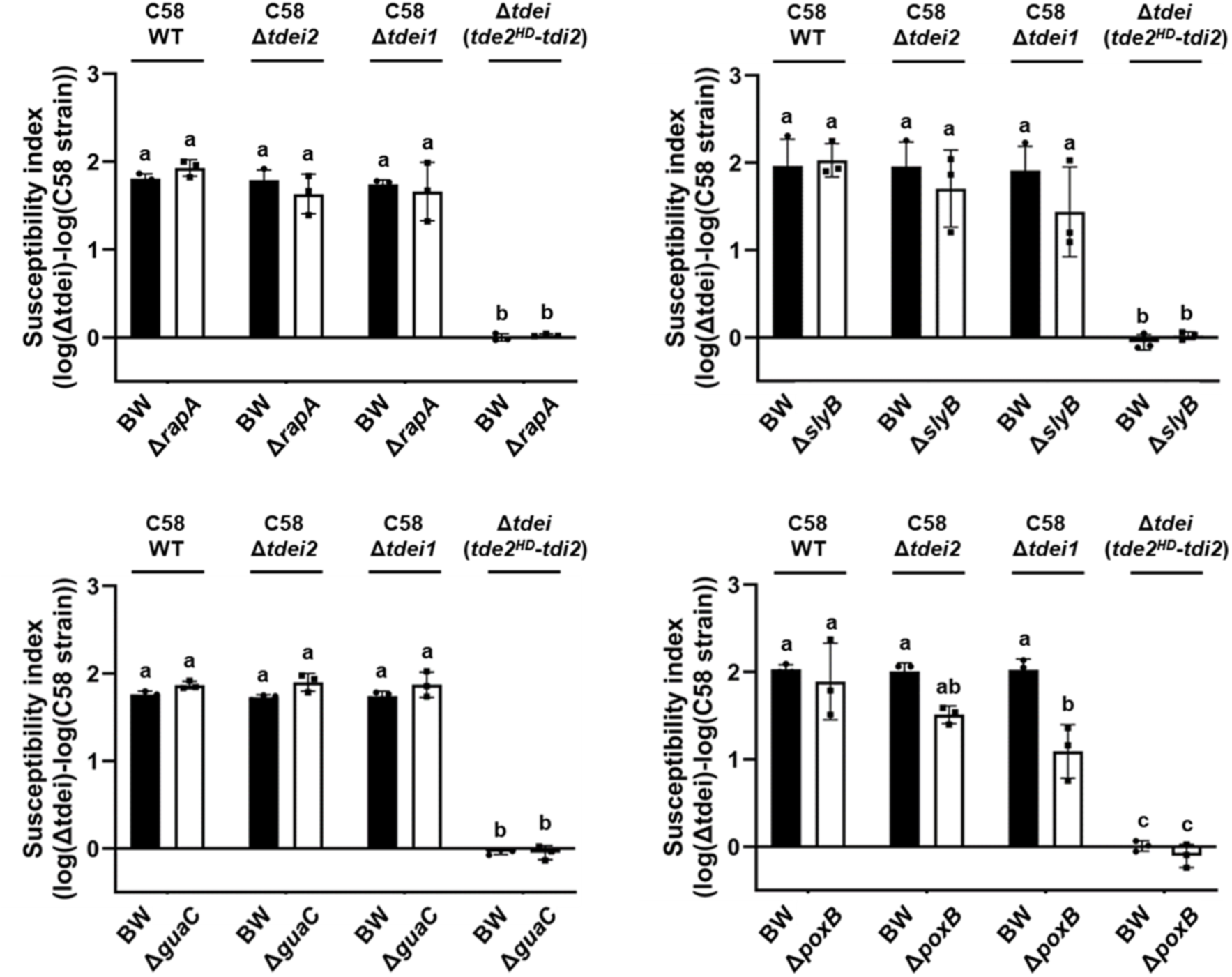
Susceptibility to Tde2-mediated interbacterial competition is reduced in the *E. coli poxB* mutant. *Agrobacterium* C58 attacker strains indicated above each plot were co-cultured with *Escherichia coli* BW25113 WT and the indicated mutant recipient strains (bottom) at a 10:1 ratio for 16 h at 25 °C. The T6SS-mediated killing effect was quantified as the susceptibility index (SI), defined as the log_10_ of the CFU count from the recipient cells co-cultured with C58 Δ*tdei* minus the log_10_ of the CFU count from the recipient cells co-cultured with other C58 strains. The dataare mean ± SD from three biological replicates. Statistical significance was determined by two-way analysis of variance (ANOVA) followed by Tukey’s honestly significant difference (HSD) test (*p* < 0.05); groups with different letters differ significantly. Susceptibility was compared among *E. coli* BW25113 WT (BW WT) and Δ*rapA*::Kan^R^ (Δ*rapA*), Δ*slyB*::Kan^R^ (Δ*slyB*), Δ*guaC*::Kan^R^ (Δ*guaC*) and Δ*poxB*::Kan^R^ (Δ*poxB*), each carrying pTrc200-HA.

**Figure 5.**
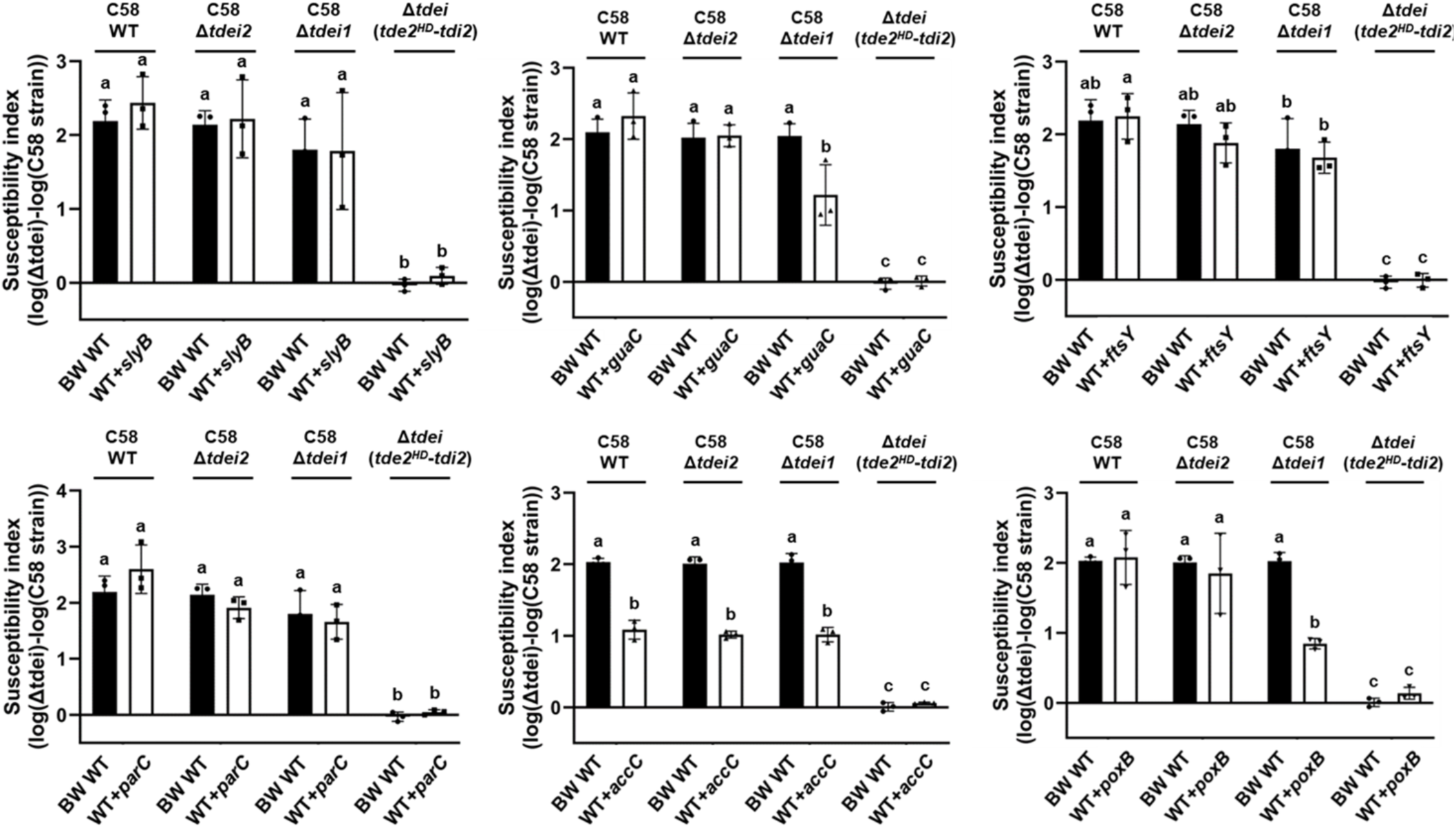
Susceptibility to Tde2-mediated interbacterial competition is reduced by overexpression of GuaC, AccC or PoxB in recipient cells. *Agrobacterium* C58 attacker strains indicated above each plot were co-cultured with *Escherichia coli* BW25113 WT and the indicated mutant recipient strains (bottom) at a 10:1 ratio for 16 h at 25 °C. The T6SS-mediated killing effect was quantified as the susceptibility index (SI), defined as the log_10_ of the CFU count from the recipient cells co-cultured with C58 Δ*tdei* minus the log_10_ of the CFU count from the recipient cells co-cultured with other C58 strains. The data are mean ± SD from three biological replicates. Statistical significance was determined by two-way analysis of variance (ANOVA) followed by Tukey’s honestly significant difference (HSD) test (*p* < 0.05); groups with different letters differ significantly. Susceptibility was compared among *E. coli* BW25113 WT carrying pTrc200-HA (BW WT) and WT overexpressing SlyB (WT+*slyB*), GuaC (WT+*guaC*), FtsY (WT+*ftsY*), ParC (WT+*parC*), AccC (WT+*accC*) and PoxB (WT+*poxB*).

### GuaC catalytic site variant remains inhibitory and associates with the Tde2 DNase domain

We therefore focused on GuaC, which specifically reduces Tde2-mediated interbacterial competition. GuaC is a guanosine 5’-monophosphate (GMP) reductase that catalyzes the irreversible, NADPH-dependent conversion of GMP to inosine monophosphate (IMP) (Kessler and Gots, 1985). To test whether GuaC enzymatic activity contributes to the reduction of Tde2-mediated interbacterial competition, we generated a catalytic site variant of GuaC (GuaC^C186A^) (Li *et al*., 2006) and expressed it from the IPTG-inducible *trc* promoter on pTrc200-HA for interbacterial competition assays. C58 WT and Tde1-mediated Δ*tdei2* attackers showed similar killing of all recipient strains (Fig. 6). By contrast, the killing outcome of the Δ*tdei1* attacker was reduced against recipient strains overexpressing WT GuaC (WT+*guaC*), and the Δ*guaC* expressing WT GuaC (*guaC^+^*) or the catalytic variant (C186A*^+^*) relative to BW WT (Fig. 6). These results suggest that GuaC enzymatic activity may be dispensable for inhibition of Tde2-mediated interbacterial competition.

**Figure 6.**
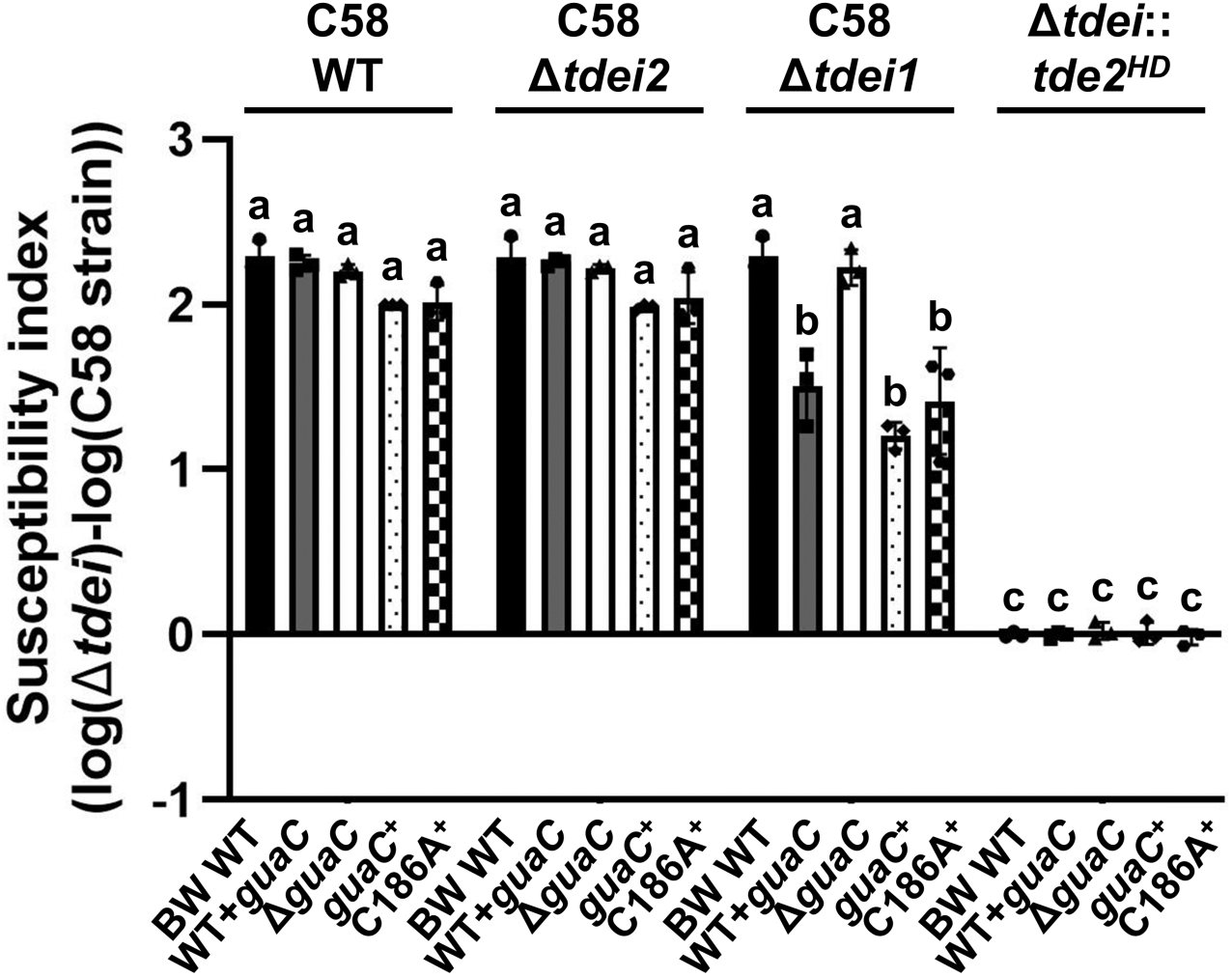
GuaC catalytic site variant remains inhibitory to Tde2-mediated interbacterial competition. *Agrobacterium* C58 attacker strains indicated above each plot were co-cultured with *Escherichia coli* BW25113 WT and the indicated mutant recipient strains (bottom) at a 10:1 ratio for 16 h at 25 °C. The T6SS-mediated killing effect was quantified as the susceptibility index (SI), defined as the log_10_ of the CFU count from the recipient cells co-cultured with C58 Δ*tdei* minus the log_10_ of the CFU count from the recipient cells co-cultured with other C58 strains. The data are mean ± SD from three biological replicates. Statistical significance was determined by two-way analysis of variance (ANOVA) followed by Tukey’s honestly significant difference (HSD) test (*p* < 0.05); groups with different letters differ significantly. Susceptibility was compared among *E. coli* BW25113 WT carrying pTrc200-HA (BW WT), WT overexpressing GuaC (WT+*guaC*), Δ*guaC*::Kan^R^ carrying pTrc200-HA (Δ*guaC*), and Δ*guaC*::Kan^R^ complemented with GuaC (*guaC^+^*) or GuaC^C186A^ (C186A^+^).

We next dissect the region of Tde2 involved in interacting with GuaC. To this end, we generated a His-tagged Tde2 DNase-domain construct (Tde2^261-526^), lacking the N-terminal region (Fig. 7A), and co-expressed HA-tagged GuaC with His-tagged WT Tde2, Tde2^HD^ or Tde2^261-526^ in the Δ*guaC*Δ*clpS* double mutant. Ni-NTA pull-down assays showed that GuaC-HA was co-purified only with the proteins containing the WT DNase domain (Tde2 and Tde2^261-526^), but not with the DNase-deficient variant (His-Tde2^HD^) or the His-sfGFP control (Fig. 7B). Ectopic expression of His-Tde2^261-526^ caused only slight growth inhibition in BW25113 WT and *clpS* mutant relative to the empty vector control and full-length WT Tde2, suggesting this partial toxicity was not dependent on ClpS (Fig. 7C). Together, these results suggest that although full-length Tde2 is required for full toxicity, the DNase domain alone is sufficient for specific association with GuaC, and this association depends on an intact HxxD motif.

**Figure 7.**
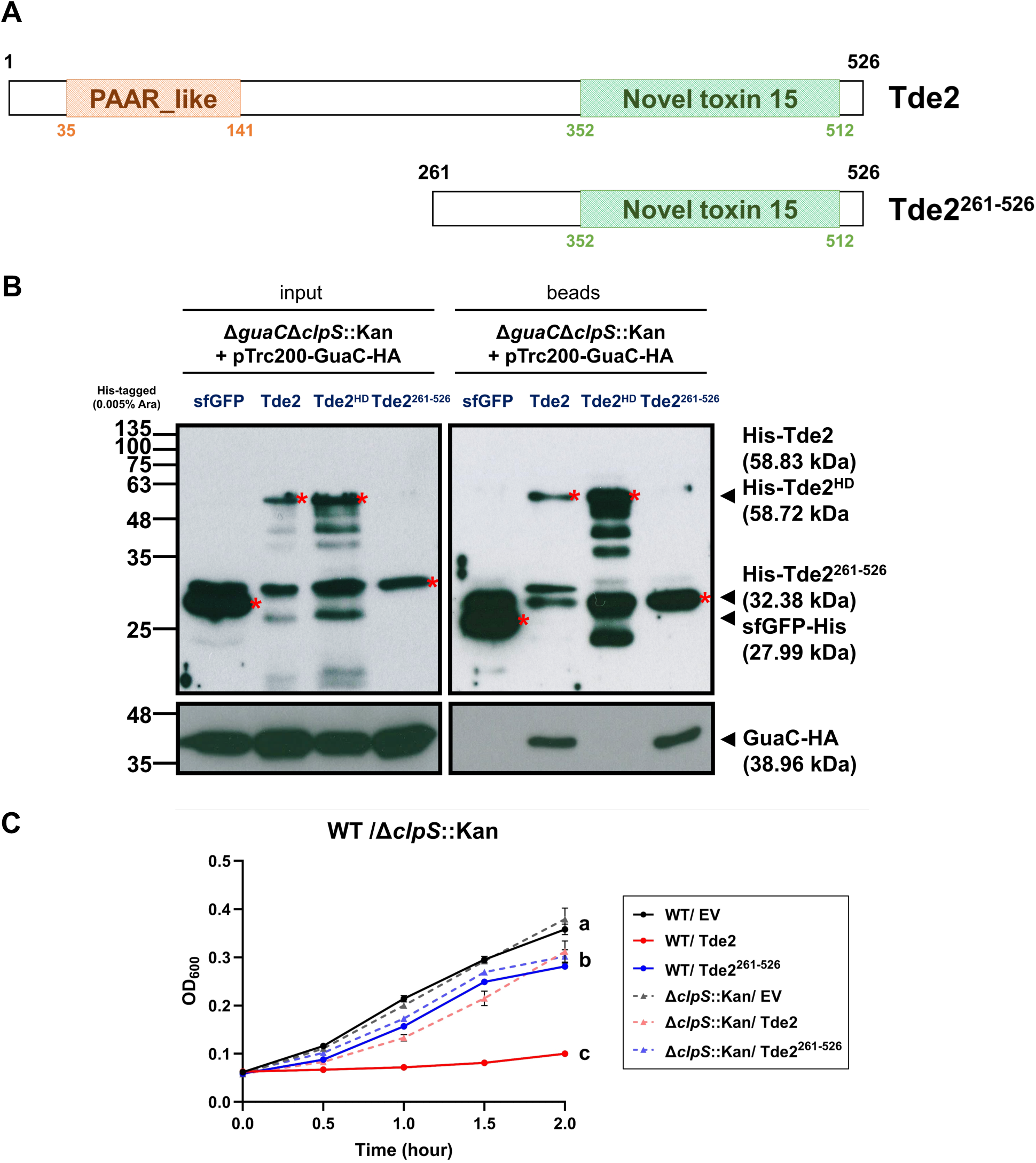
The Tde2 DNase domain is sufficient for GuaC interaction and exhibits weak ClpS-dependent toxicity. **A.** Schematic of the predicted domain organization of Tde2 based on Basic Local Alignment Search Tool (BLAST) analyses. Tde2 contains an N-terminal PAAR-like domain (residues 35–141) and a C-terminal Ntox15 DNase domain (residues 352–512). The truncated variant Tde2^261-526^ comprises the Ntox15 DNase domain. **B.** His-tagged Tde2, Tde2^HD^, Tde2^261-526^ or sfGFP were co-expressed with HA-tagged GuaC in Δ*guaC*Δ*clpS* double mutant and induced with 0.005% (w/v) L-arabinose for 2 h. Cells were lysed, and His-tagged proteins together with their associated partners were co-precipitated using Ni-NTA agarose beads. Protein samples were analyzed by immunoblotting with anti-His and anti-HA antibodies. Input: Supernatant from cell lysates before purification. Beads: Ni-NTA agarose bead-bound fractions containing His-tagged proteins and their associated proteins. Asterisks (*) indicate the expected protein bands. **C.** Growth of *Escherichia coli* BW25113 WT (solid line) and *clpS* mutant (dashed line) carrying empty vector (EV, black) or plasmid expressing His-tagged Tde2 (red) or the truncated variant Tde2^261-526^ (blue). Cultures were supplemented with 0.005% L-arabinose at 0 h. The data represent mean ± SD from four biological replicates. Statistical significance was determined by one-way analysis of variance (ANOVA) followed by Tukey’s honestly significant difference (HSD) test (*p* < 0.05); groups with different letters differ significantly.

## Discussion

Our study identifies the recipient ClpAPS protease system as a determinant of susceptibility to T6SS-mediated intoxication by the *Agrobacterium* DNase effector Tde2. Across interbacterial competition assays and ectopic expression experiments, efficient Tde2-mediated toxicity required recipient ClpAPS, indicating that a conserved recipient proteostasis pathway promotes full intoxication. Based on these findings, we propose a working model in which ClpAPS enhances Tde2 activity indirectly, potentially by clearing recipient factors that otherwise limit its DNase function (Fig. 8). Together, our results identify the ClpAPS protease system as an unexpected recipient susceptibility (RS) factor that shapes the outcome of T6SS-mediated competition across multiple Gram-negative bacterial contexts.

**Figure 8.**
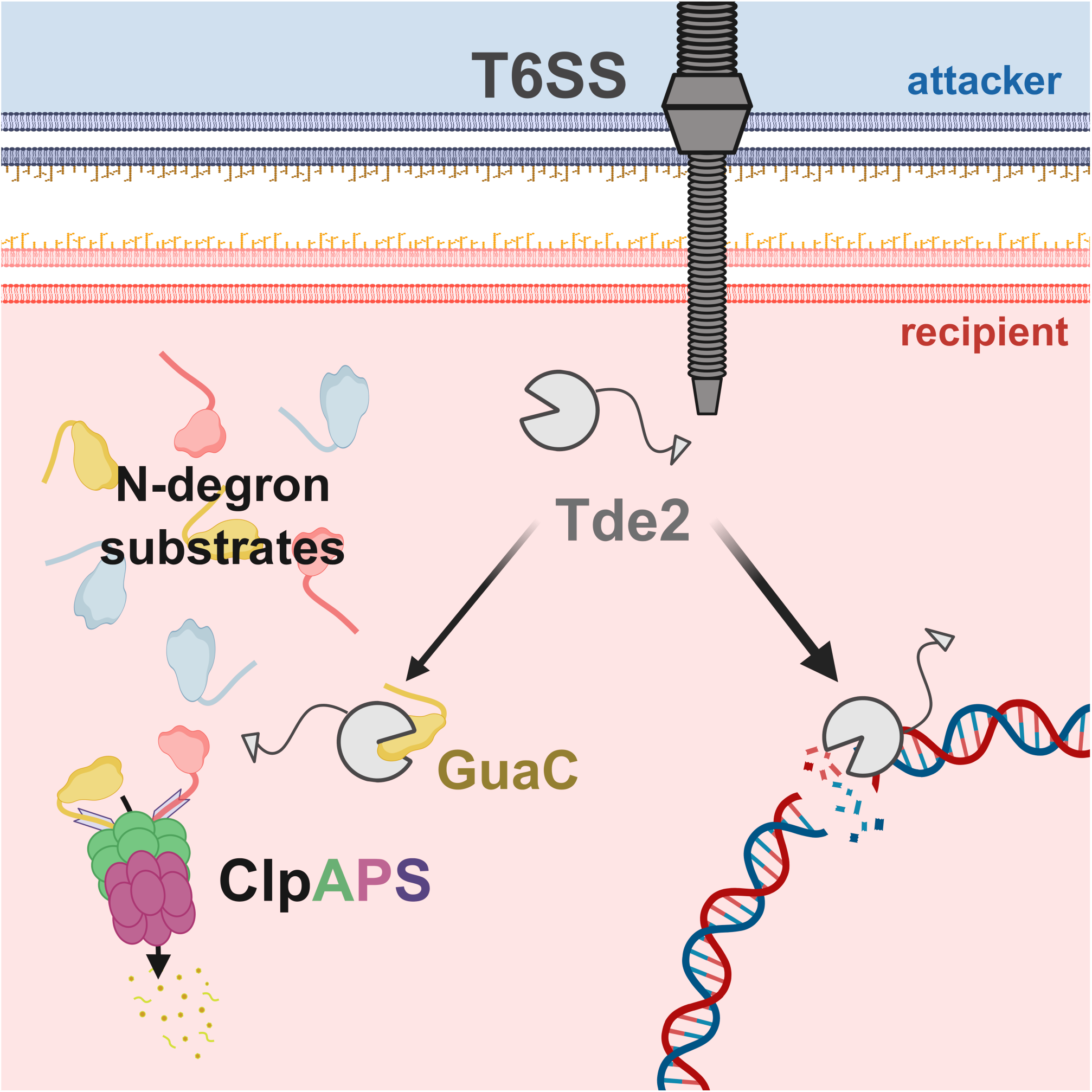
Proposed model for indirect activation of Tde2 by the recipient ClpAPS protease complex. Upon *Agrobacterium* T6SS attack, the DNase effector Tde2 is delivered into recipient cells, where it induces DNA damage. Tde2-mediated toxicity depends indirectly on recipient ClpAPS through clearance of recipient factors that otherwise limit its DNase function. These factors may include candidate N-degron substrates, such as GuaC, that associate with Tde2. When ClpAPS expression or activity is reduced, these factors persist, Tde2 activity is reduced, and DNA damage is diminished. Illustration created in BioRender.

RS factors have emerged as important modulators of T6SS-mediated competition. In addition to cognate immunity systems and inducible defence responses, recipient cells can encode functions that are exploited by incoming effectors or that otherwise influence competition outcomes (Hersch *et al*., 2020; Lin *et al*., 2020a; Robitaille *et al*., 2021). Many previously described RS factors act in the periplasm, where they facilitate effector folding or activation (Mariano *et al*., 2018) or contribute to cell envelope modification and maintenance (Trotta *et al*., 2023; Tsai *et al*., 2025). By contrast, our findings support a cytoplasmic mechanism in which recipient proteostasis indirectly modulates effector activity after delivery.

The requirement for ClpAPS in recipient *E. coli*, *Agrobacterium*, and *D. dadantii* during Tde2-mediated interbacterial competition suggests that this pathway may have broad relevance across Gram-negative bacteria. Consistent with this view, *Vibrio cholerae* T6SS-mediated competition also depends on recipient *E. coli* ClpP (Hersch *et al*., 2021), although the underlying mechanism remains elusive. Because *V. cholerae* does not encode a Tde-like DNase effector (Crisan and Hammer, 2020), ClpP-dependent susceptibility in this system is likely to rely on a mechanism distinct from that proposed here. Given the conserved role of ClpAPS in degrading N-degron substrates (Katayama-Fujimura *et al*., 1987; Dougan *et al*., 2002; Wong and Houry, 2004), we provide evidence that some of the Tde2-associated N-degron substrates identified in the absence of ClpP could modulate Tde2 toxicity (Fig. 4, 5). Among them, GuaC emerged as the strongest mechanistic lead, as its overexpression selectively reduced Tde2-mediated competition. These findings are consistent with the idea that at least some recipient factors can interact directly with Tde2 and modulate its activity. Defining how such factors interact with Tde2 will be important for understanding how recipient cytoplasmic proteostasis influences effector activity. Consistent with a possible reciprocal relationship between Tde2 and the ClpAPS pathway, we also observed that ClpA abundance was reduced upon expression of active Tde2 in *E. coli* strains carrying ClpP (Fig. 2B). Although the basis and physiological relevance of this effect remain unclear, this finding raises the possibility that Tde2 not only depends on recipient ClpAPS activity but may also perturb this pathway during intoxication. It is intriguing that Tde2 interacts with GuaC in a manner dependent on an intact DNase domain.

This observation suggests that GuaC may bind to Tde2 via the HxxD catalytic motif, or alternatively, that it preferentially recognizes the active form of Tde2. Notably, Tde2^261-526^ remains sufficient for interaction with GuaC despite exhibiting only weak DNase toxicity (Fig. 7), indicating that GuaC binding is not strictly coupled to full catalytic activity. Based on these findings, we propose that GuaC interacts with the HxxD motif of Tde2, thereby attenuating its DNase activity. Although both Tde1 and Tde2 belong to the Ntox15 superfamily and share a conserved HxxD catalytic motif required for DNase activity (Ma *et al*., 2014), GuaC selectively affected Tde2-mediated interbacterial competition. Due to the instability of Tde1 during heterologous expression in *E. coli* (Ali *et al*., 2023), we were unable to obtain sufficient protein to assess whether GuaC can also recognize Tde1; the basis of this specificity remains unresolved. Nevertheless, the marked divergence between the C-terminal DNase domains of Tde1 and Tde2, which share only 29.14% identity (49% similarity) (Fig. S4). Therefore, we speculate that recognition by GuaC, along with other N-degron substrates that inhibit Tde2 DNase toxicity, depends on features unique to Tde2 rather than on the shared Ntox15 fold alone.

In conclusion, our findings support a model in which recipient ClpAPS enhances Tde2 intoxication by counteracting cytoplasmic constraints on its DNase activity. Rather than acting directly on Tde2, recipient ClpAPS appears to promote a cytoplasmic state that is permissive for Tde2-mediated DNA damage. Because ClpAPS carries out a conserved housekeeping role in protein quality control (Dougan *et al*., 2002; Zeth *et al*., 2002), it may provide a stable recipient function that Tde2 can exploit across diverse bacterial species. This dependence may in turn reduce the selective pressure for Tde2 to evolve complete independence from the ClpAPS pathway. The findings also expand the concept of recipient susceptibility by showing that conserved cellular quality-control pathways can potentiate the activity of a delivered T6SS effector. More broadly, they highlight that the outcome of interbacterial competition is shaped not only by effector repertoires and defence systems, but also by the physiological state of the target cell. As T6SS continues to emerge as a key determinant of bacterial fitness and niche dominance (Ho *et al*., 2014; Coulthurst, 2019; Mijatović Scouten *et al*., 2025), elucidating the molecular basis of RS factors will provide important insight into how bacterial species balance competition, survival, and community assembly within complex microbial communities.

## Materials and Methods

### Bacterial Strains and Plasmids

The bacterial strains and plasmids used in this study are listed in Table S1. *Escherichia coli* and *Dickeya dadantii* strains were cultured in Luria-Bertani (LB) medium supplemented with appropriate antibiotics at 37°C with shaking at 220 rpm, whereas *Agrobacterium* strains were grown in 523 medium (Kado and Heskett, 1970) at 25°C with shaking at 220 rpm. The *E. coli* BW25113 WT strain and its Keio knockout mutants (Baba *et al*., 2006) were obtained from the Keio collection (NBRP, NIG, Japan). Plasmids were maintained in *E. coli*, *Agrobacterium* and *Dickeya dadantii* using the following antibiotic concentrations: kanamycin (20 μg/ml), spectinomycin (100 μg/ml) and gentamicin (30 μg/ml).

*Agrobacterium* mutants were generated by double-crossover homologous recombination using the suicide plasmid pJQ200ks (Table S1), which carries the *sacB* and gentamicin resistance genes for selection, as described previously (Ma *et al*., 2009). Mutants were selected based on sucrose resistance and antibiotic sensitivity. *E. coli* double-deletion mutant was constructed using the λ Red recombinase–mediated recombination system, as previously described (Datsenko *et al*., 2000). Briefly, target genes were replaced with antibiotic resistance cassettes by homologous recombination. All mutations were confirmed by polymerase chain reaction (PCR) and Sanger sequencing.

For plasmid construction, target genes were amplified by PCR using gene-specific primers (Table S2) and cloned into the indicated vectors using restriction enzyme digestion and T4 DNA ligase–mediated ligation. Recombinant plasmids were transformed into *E. coli* DH5α, and positive clones were verified by Sanger sequencing. Verified plasmids were subsequently introduced into the indicated bacterial strains by PEG-mediated chemical transformation (Chung *et al*., 1989) or electroporation, as appropriate.

### Antibody generation

Polyclonal antibodies against ClpP and ClpA were generated in rabbits by immunization with purified His-tagged ClpP and His-tagged ClpA proteins, respectively. Antibodies against ClpS and Tde2 were generated by rabbit immunization with synthetic peptides corresponding to residues 24-42 of ClpS (PPSMYKVILVNDDYTPMEF-Cys) and to the C-terminus of Tde2 (CPSNPSRPGEPLTGT and SRLTEHAKRQAANNC). Peptide synthesis, animal immunization, and serum collection were performed by Yao-Hong Biotech Inc. (Taiwan).

### Interbacterial competition assay

Overnight cultures of *Agrobacterium* C58 strains (attacker) and recipient strains (*E. coli* BW25113, *D. dadantii* 3937, or *Agrobacterium* C58 Δ3TIS) harboring pTrc200-HA or its derivatives were harvested, washed with 0.9% (w/v) NaCl and adjusted to OD_600_ = 1.0 (attackers) and 0.1 (recipients), respectively. Two 10-µl aliquots of the mixed bacterial suspension were spotted onto *Agrobacterium* Kill-triggering (AK) medium solidified with 2% (w/v) agar (Lin *et al*., 2020b; Yu *et al*., 2021), air-dried for 40 min and incubated at 25°C for 16 h. After incubation, the bacterial spots were resuspended in 1 ml of 0.9% NaCl, serially diluted and plated onto LB agar supplemented with spectinomycin to recover recipient cells. Plates were incubated at 37°C for *E. coli* and *D. dadantii*, or 25°C for *Agrobacterium*. Colony-forming units (CFUs) were enumerated, and the competition outcome was quantified using the susceptibility index (SI) defined as the logarithm of the recipient CFUs recovered after co-culture with C58 Δ*tdei* subtracted by that recovered after co-culture with other C58 strains (Lin *et al*., 2020b). The data are presented as mean ± standard error (SE) from three independent biological replicates. Statistical analysis was determined using Tukey’s honestly significant difference (HSD) test.

### Growth inhibition assay

Growth inhibition assays were performed as described (Ma *et al*., 2014; Ali *et al*., 2023) with minor modifications. Freshly transformed *E. coli* BW25113 strains carrying pJN105 or its Tde2 variants were grown overnight in LB with gentamicin, diluted to OD_600_ = 0.1 in induction medium (LB containing gentamicin and 0.005% [w/v] L-arabinose), and cultured in 96-well plates (200 µl per well). OD_600_ was recorded every 15 min for 2 h at 37°C with shaking (237 cpm) using a Synergy H1 Multimode Reader (BioTek, United States). The data represent the mean ± SD of six (or four) biological replicates from three independent experiments. Statistical significance was determined by one-way ANOVA (*p* < 0.001) followed by Scheffé’s test.

### Sodium dodecyl sulfate–polyacrylamide (SDS-PAGE) gel electrophoresis and Western blot analysis

The bacterial cells were adjusted to OD_600_ = 2 or 10, pelleted (16,000 × g, 10 minutes), and resuspended in 10 mM Tris-HCl (pH 8.5) with 2× SDS sample loading buffer. Samples were boiled at 95°C for 10 minutes, separated by SDS-PAGE, and transferred to Immobilon-P membranes (Merck Millipore, United States). Primary antibodies were used at the following dilutions: polyclonal anti-Tde2 (1: 8,000; Yao-Hong Biotech Inc., Taiwan), polyclonal anti-ClpP (1: 20,000; Yao-Hong Biotech Inc., Taiwan), polyclonal anti-ClpA (1: 10,000; Yao-Hong Biotech Inc., Taiwan), polyclonal anti-ClpS (1: 2,000; Yao-Hong Biotech Inc., Taiwan), polyclonal anti-EF-Tu (1: 6,000; Dr. Chu, Hsiu-An’s Lab, Taiwan), monoclonal anti-His (1: 10000; Merck Millipore, United States), monoclonal anti-HA (1: 10000; Sigma-Aldrich, United States). HRP-conjugated secondary antibodies (goat anti-rabbit or rabbit anti-mouse; 1: 10,000; GeneTex, Taiwan) were applied, and signals were detected using Western Lightning ECL Pro (PerkinElmer Life Sciences, United States) and visualized on X-ray film using an automatic film processor.

### Plasmid DNA degradation assay

Plasmid DNA degradation assays were performed as described for the **Growth inhibition assay** with minor modifications. The *E. coli* cultures were induced with 0.005% (w/v) L-arabinose for 1 h, collected, and normalized to OD_600_ = 10. Plasmid DNA was extracted and analyzed by agarose gel electrophoresis. All experiments were performed in three independent biological replicates.

### Terminal deoxynucleotidyl transferase dUTP nick end labeling (TUNEL) assay analysis

TUNEL assays were also performed as described for the **Growth inhibition assay** with minor modifications. The *E. coli* cultures were induced with 0.005% (w/v) L-arabinose for 1 h, collected, and normalized to OD_600_ = 1.0. Bacterial cells were fixed with 1% (w/v) paraformaldehyde in PBS (pH 7.4) for 40 min on ice and stained using the APO-DIRECT™ Kit (BD Biosciences, United States) according to the manufacturer’s instructions. Samples were incubated at 28°C for 20 h, and analyzed using a CytoFLEX S flow cytometer (Beckman Coulter, United States). All experiments were performed in three independent biological replicates.

### Pull-down assay

His-tagged pull-down assays by Ni-NTA were carried out according to the manufacturer’s instructions (Cytiva, Switzerland). His-tagged proteins (Tde2 variants and sfGFP) were expressed from pJN105 in *E. coli* and induced with 0.005% (w/v) L-arabinose. Bacterial cells were harvested and resuspended in lysis buffer (50 mM NaH_2_PO_4_, 300 mM NaCl, 10 mM imidazole, 1 mg /ml lysozyme, 40 μl/ml 25X EDTA-free protease inhibitor (Roche, Switzerland), 0.1 mg/ml DNase I and 0.1 mg/ml RNase A, pH 8.0) to OD_600_ = 4.0. Two milliliters of bacterial cells were lysed using a Continuous Flow Cell Disruptor (TS Series 0.75kw Model, Constant Systems Ltd., United Kingdom), and the supernatant was collected by centrifugation at 18,000 × g for 1 h. The cleared lysate was incubated with Ni-NTA agarose beads (Ni Sepharose^TM^ 6 Flat Flow, Cytiva, Switzerland) overnight at 4°C. Two hundred microliters of beads was washed with wash buffer (50 mM NaH_2_PO_4_, 300 mM NaCl, 20 mM imidazole, 40 μl/ml EDTA-free protease inhibitor, pH 8.0). His-tagged proteins (Tde2 variants and sfGFP) and their associated partners were either eluted with 2× SDS sample loading buffer for Western blot analysis or processed for liquid chromatography–tandem mass spectrometry (LC–MS/MS).

### Liquid chromatography-tandem mass spectrometry (LC-MS/MS) sample preparation and analysis

LC-MS/MS sample preparation followed a previously described protocol (Chen *et al*., 2023), except that phosphopeptide enrichment was omitted. Two independent experiments were performed. In the first experiment, samples were analyzed on a Q Exactive Orbitrap mass spectrometer coupled to an UltiMate 3000 RSLCnano system and Nanospray Flex ion source (Thermo Fisher Scientific, United States). Peptides were separated on a nanoEase C18 capillary column (130 Å, 1.7 μm, 75 μm × 250 mm; Waters, United States) using a binary solvent system consisting of buffer A (0.1% formic acid in water) and buffer B (0.1% formic acid in acetonitrile). A linear gradient from 3% to 25% buffer B over 60 min was followed by an increase to 85% buffer B over 2 min, column washing, and re-equilibration at 3% buffer B at 450 nL/min. The instrument was operated in the data-dependent mode (Full MS/ddMS², TopN). Full MS scans (m/z 350-1,600) were collected at 70,000 resolution (FWHM) with an automatic gain control (AGC) target of 3 × 10^6^. MS/MS spectra were acquired by higher-energy collisional dissociation (HCD) at 27% normalized collision energy (NCE), 17,500 resolution, a maximum injection time of 100 ms, and an AGC target of 1 × 10^5^. The top 10 precursor ions were selected for fragmentation using a 2.0 m/z isolation window and a dynamic exclusion time of 20 sec.

Raw data were processed in Proteome Discoverer™ v2.4.0.305 (Thermo Fisher Scientific, United States) using the SEQUEST HT and Mascot search algorithms against the *E. coli* BW25113 protein database. High-confidence proteins were defined at *q* ≤ 0.05 (5% false discovery rate, FDR). Identified proteins were analyzed using Venny 2.1 to determine the known N-degron substrates (ClpS-binding proteins) among Tde2-associated proteins, and Venn diagrams were generated comparing proteins associated with His-tagged Tde2, sfGFP, and the empty vector (EV) controls.

In the second experiment, samples were loaded onto Evotip Pure and analyzed using an Evosep One liquid chromatography system (Evosep Biosystems, Denmark) coupled to a timsTOF HT mass spectrometer (Bruker Daltonics, United States). Peptides were separated using the Extended method on an EV1137 performance column (15 cm × 150 μm ID, 1.5 μm; Evosep Biosystems, Denmark) maintained at 40°C within a CaptiveSpray ion source (Bruker Daltonics, United States). The instrument was operated in data-dependent acquisition parallel accumulation-serial fragmentation (dda-PASEF) mode with 10 PASEF MS/MS scans per TopN acquisition cycle. MS/MS spectra were acquired over m/z 100-1700 and an ion mobility range of 0.6-1.6 Vs/cm^2^, with 100 ms accumulation and ramp times. Collision energy was applied as a linear ramp from 20 eV at 1/K_0_ = 0.6 Vs/cm^2^ to 59 eV at 1/K_0_ = 1.6 Vs/cm^2^ Vs/cm^2^. Singly charged precursor ions were excluded using a polygon filter in the m/z-ion mobility plane. Precursors with intensities above 2,500 were selected for fragmentation using isolation windows of 2 Th (m/z < 700) and 3 Th (m/z > 800), and precursors reaching a target intensity of 20,000 were dynamically excluded for 24 sec.

Raw data were processed using SpectroMine™ v4.5.240625.52329 (Biognosys, Switzerland) with standard parameters. High-confidence identifications were defined at *q* ≤ 0.05 (5% FDR) and applied at the peptide-spectrum match (PSM), peptide, and protein levels, allowing up to two missed cleavages. Proteins detected in at least two of four biological replicates were subjected to fold-change analysis, comparing proteins associated with His-Tde2 to negative controls (EV and sfGFP-His). Enrichment of known N-degron substrates (ClpS-binding proteins) among Tde2-associated proteins was defined using a ≥ 2-fold-change threshold in both comparisons (His-Tde2 vs EV and His-Tde2 vs sfGFP-His).

## Acknowledgements

The authors acknowledge the Lai lab members for inspiring discussion and suggestions. We thank Yi-Chieh Wang for technical assistance on constructing the *Agrobacterium clpP* mutants and Dr. Ching-Hong Yang (University of Wisconsin-Milwaukee) for providing *Dickeya dadantii* strains. We also acknowledge the Proteomics Core Laboratory, Comprehensive Flow Cytometry Laboratory, and the Genomic Technology Core, located at the Institute of Plant and Microbial Biology, Academia Sinica, for technical assistance with LC-MS/MS analysis, TUNEL assay, and Sanger sequencing, respectively. The work in the Lai lab is supported by National Science and Technology Council (NSTC) of Taiwan (113-2311-B-001-036-MY3) and Academia Sinica Investigator Award (AS-IA-107-L01) to E.-M.L. The funders had no role in study design, data collection and interpretation, or the decision to submit the work for publication.

## Author Contributions

Y.-H.W and E.-M.L. contributed to the conceptualization of the project. Y.-H.W contributed to the investigation, methodology, data curation, formal analysis, and data visualization. H.-H.L and X.-T.Z contributed to the investigation. Y.-H.W and E.-M.L. wrote the initial manuscript. H.-H.H and E.-M.L supervised the project. All authors reviewed and edited the manuscript.

## Conflict of Interest Statement

The authors declare no conflict of interest.

## Supplementary Figure legends

**Supplementary Figure 1.**
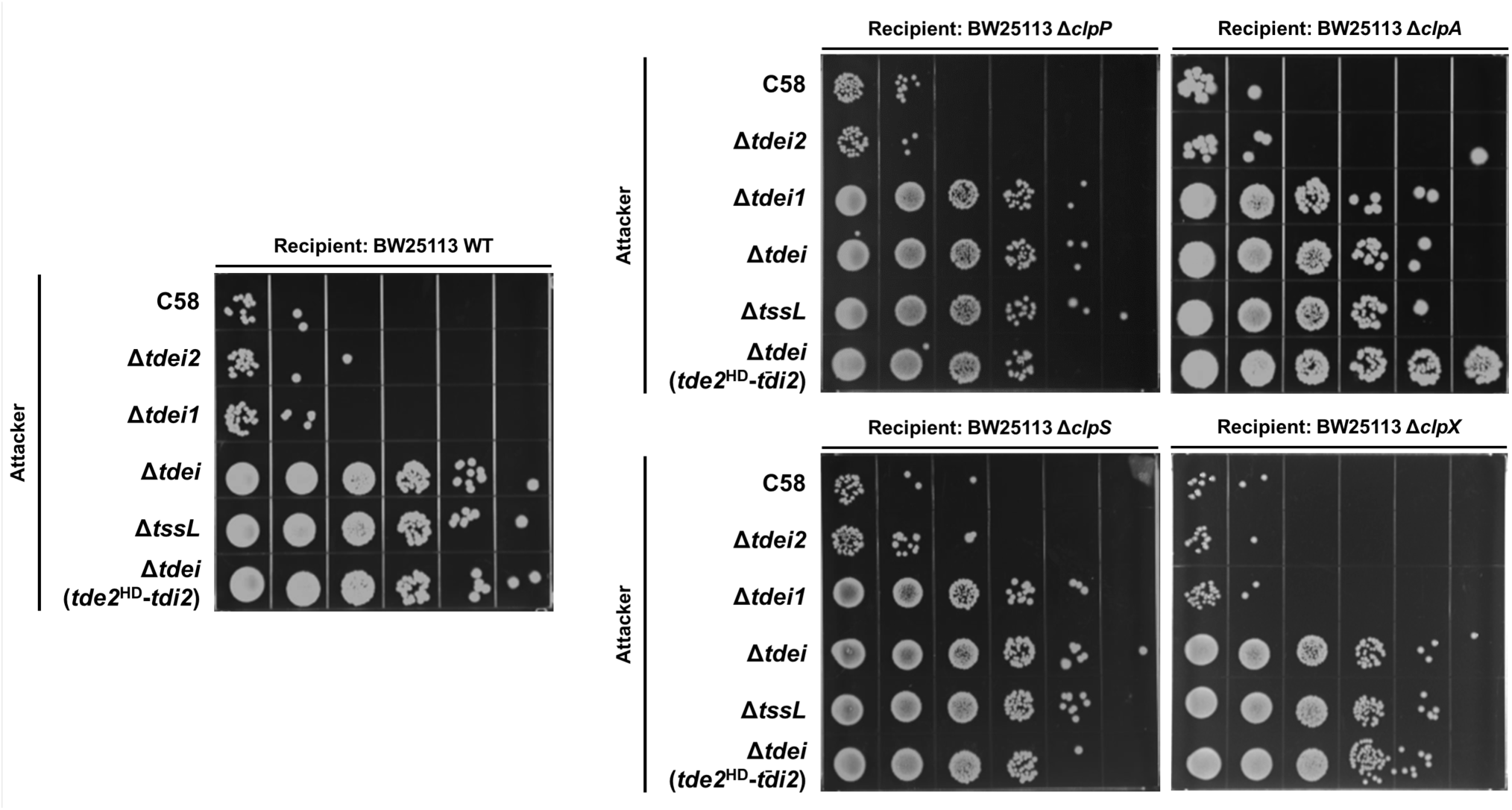
A*g*robacterium type VI secretion system (T6SS) Tde2-mediated interbacterial competition depends on the recipient ClpAPS. *Agrobacterium* C58 attacker strains indicated at left were mixed with *Escherichia coli* BW25113 recipient strains indicated above each panel, and recipient survival was assessed by spotting on agar plate with 10x serial dilution after interbacterial competition. The T6SS-defective *tssL* mutant attacker served as a negative control, and the *tdei* mutant attacker showed a comparable defect in killing. Recipient strains were BW25113 WT (BW WT), Δ*clpP*::Kan^R^ (Δ*clpP*), Δ*clpA*::Kan^R^ (Δ*clpA*), Δ*clpS*::Kan^R^ (Δ*clpS*) and Δ*clpX*::Kan^R^ (Δ*clpX*), each carrying pTrc200-HA. Extended data of Figure 1.

**Supplementary Figure 2.**
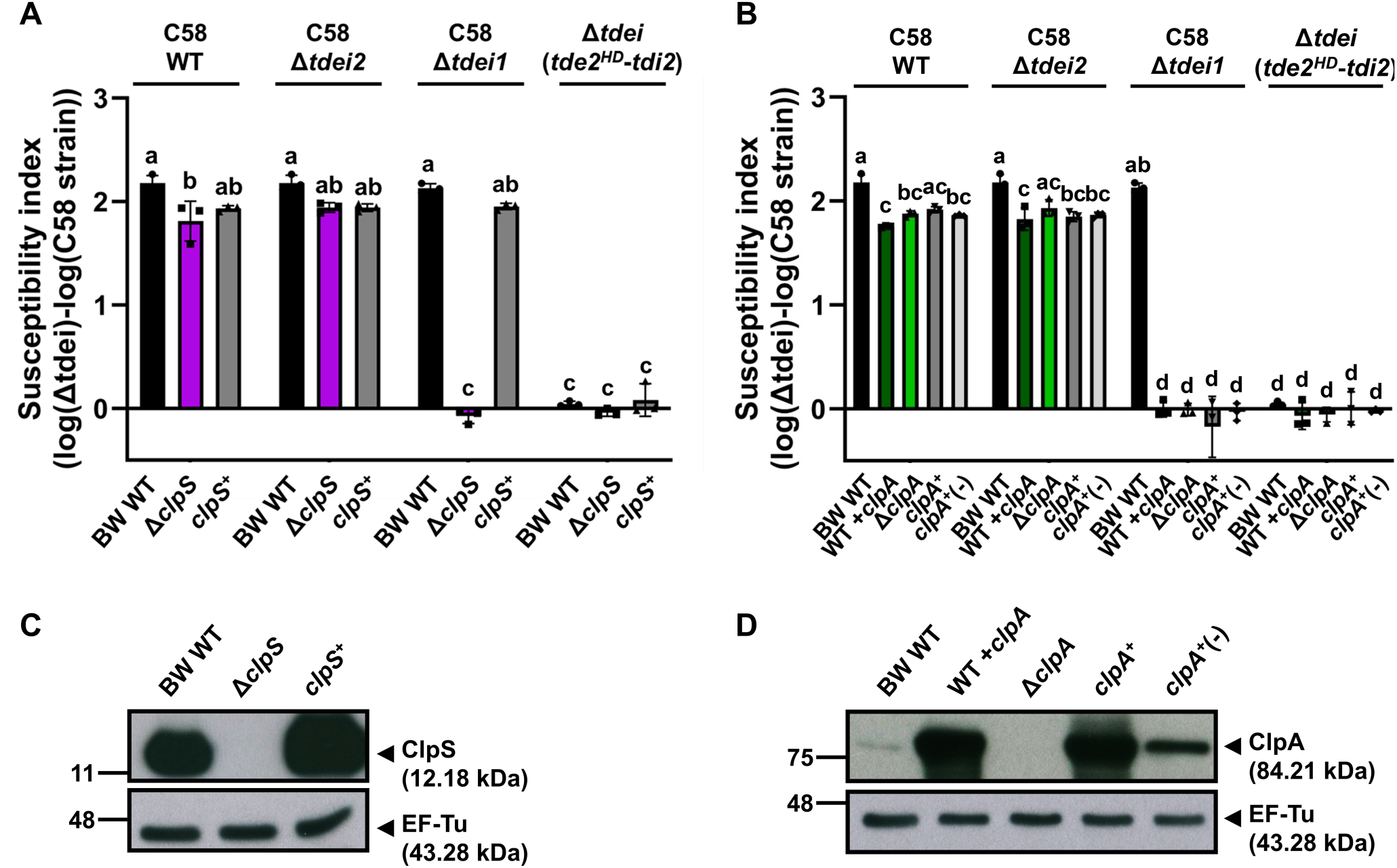
Susceptibility to *Agrobacterium* T6SS Tde2-mediated interbacterial competition is restored by ClpS, but not ClpA complementation. *Agrobacterium* C58 attacker strains indicated above each plot were co-cultured with the indicated *Escherichia coli* BW25113 recipient strains (bottom) at a 10:1 ratio for 16 h at 25 °C. The T6SS-mediated killing outcome was quantified as the susceptibility index (SI), defined as the log_10_ of the CFU count from the recipient cells co-cultured with C58 Δ*tdei* minus the log_10_ of the CFU count from the recipient cells co-cultured with other C58 strains. The data are mean ± SD from three biological replicates. Statistical significance was determined by two-way analysis of variance (ANOVA) followed by Tukey’s honestly significant difference (HSD) test (*p* < 0.05); groups with different letters differ significantly **A.** Susceptibility of *E. coli* BW25113 WT, Δ*clpS*::Kan^R^ (Δ*clpS*) and Δ*clpS* complemented strain with overexpressing ClpS (*clpS*^+^), each carrying pTrc200-HA. **B.** Susceptibility of *E. coli* BW25113 WT (BW WT), WT overexpressing ClpA (WT+*clpA*), Δ*clpA*::Kan^R^ (Δ*clpA*), Δ*clpA* complemented strain with overexpressing ClpA with (*clpA*^+^) or without (*clpA*^+^(-)) IPTG induction. **C. and D.** Immunoblot analysis of ClpA, ClpS, and Ef-Tu (loading control) from *E. coli* BW25113 recipient protein samples before the interbacterial competition. Protein markers are indicated on the left, and calculated molecular sizes of specific proteins are indicated on the right.

**Supplementary Figure 3.**
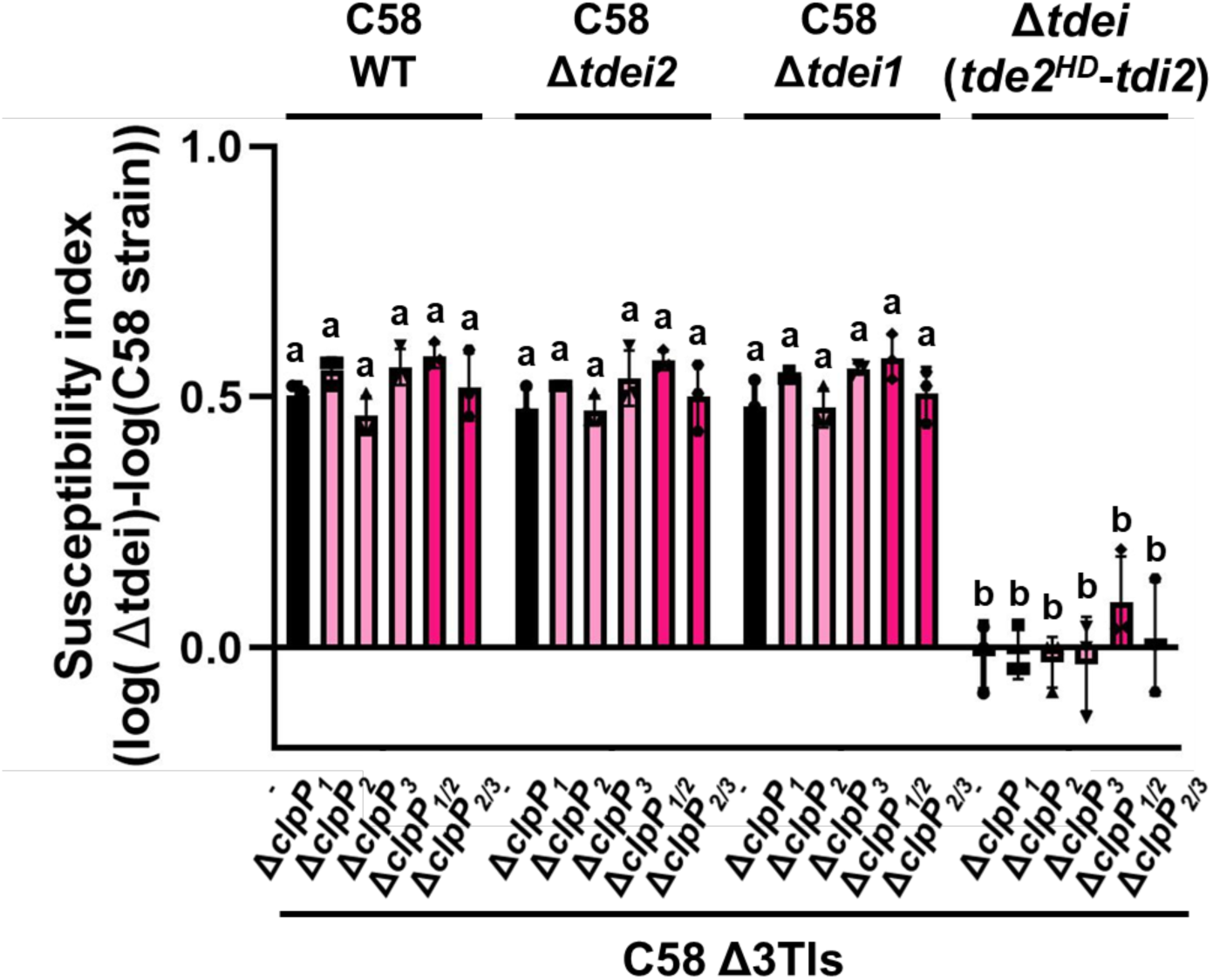
Single or double *clpP* deletions in C58 Δ3TIsdo not confer resistance to *Agrobacterium* T6SS-mediated interbacterial competition. *Agrobacterium* C58 attacker strains indicated above the plot were co-cultured with the indicated susceptible *Agrobacterium* C58 Δ3TIs recipient strains (bottom) at a 10:1 ratio for 16 h at 25 °C. The T6SS-mediated killing effect was quantified as the susceptibility index (SI), defined as the log_10_ of the CFU count from the recipient cells co-cultured with C58 Δ*tdei* minus the log_10_ of the CFU count from the recipient cells co-cultured with other C58 strains. The data are mean ± SD from three biological replicates. Statistical significance was determined by two-way analysis of variance (ANOVA) followed by Tukey’s honestly significant difference (HSD) test (*p* < 0.05); groups with different letters differ significantly. Susceptibility was compared among *Agrobacterium* C58 Δ3TIs, Δ3TIs *clpP_1_* mutant (Δ*clpP_1_*), Δ3TIs *clpP_2_* mutant (Δ*clpP_2_*), Δ3TIs *clpP_3_* mutant (Δ*clpP_3_*), Δ3TIs *clpP_1/2_* double mutant (Δ*clpP_1/2_*) and Δ3TIs *clpP_2/3_* double mutant (Δ*clpP_2/3_*) each carrying pTrc200-HA.

**Supplementary Figure 4.**
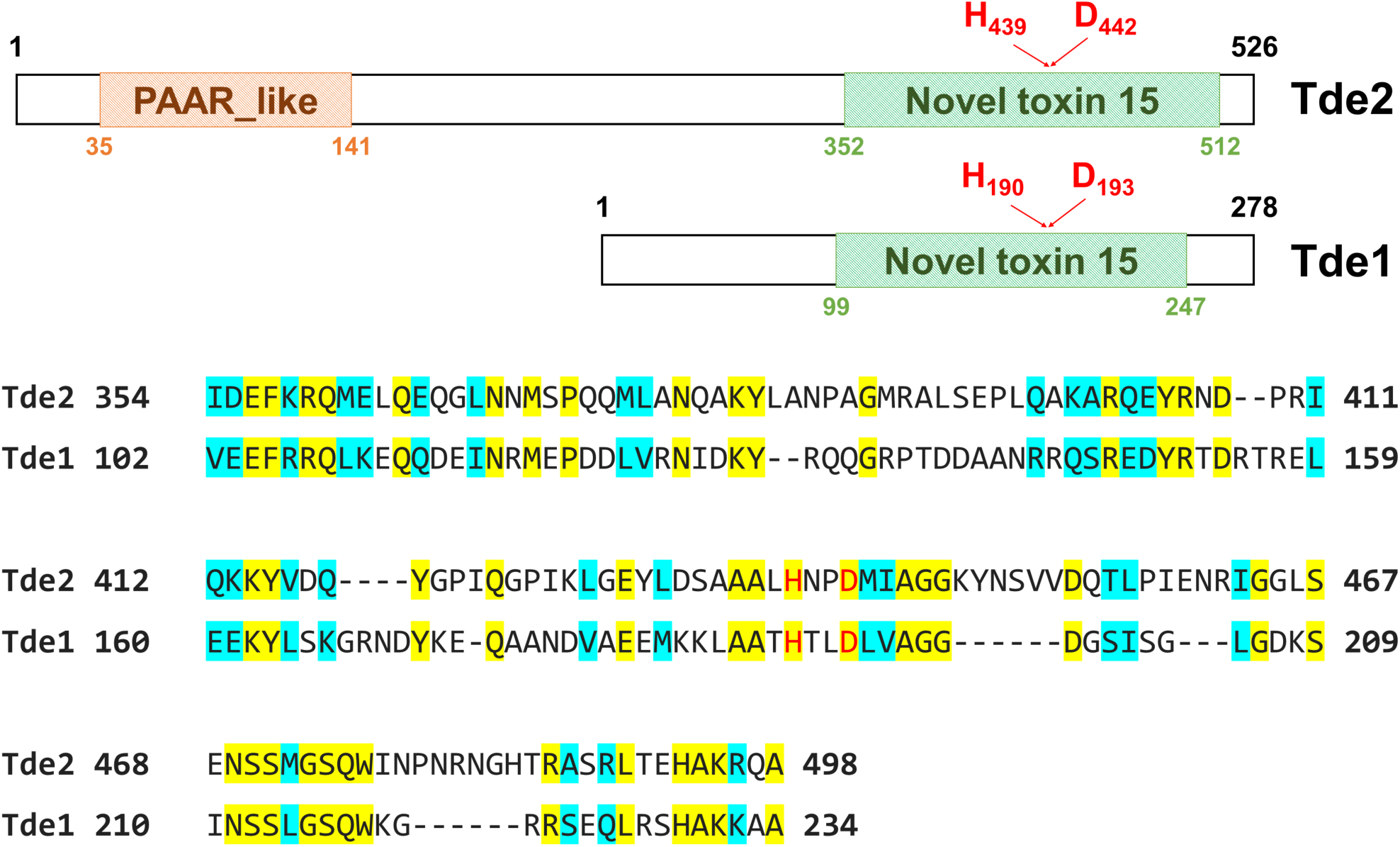
Alignment of the Ntox15 DNase domains of Tde1 and Tde2. Schematic representation of Tde2 and Tde1 domain organization based on Basic Local Alignment Search Tool (BLAST) analyses is shown above. Tde2 contains an N-terminal PAAR-like domain and a C-terminal Ntox15 domain, whereas Tde1 contains a single Ntox15 domain. Amino acid sequence alignment of the Ntox15 regions of Tde2 and Tde1 is shown below. Identical residues are highlighted in yellow, conserved residues are highlighted in cyan, and the predicted catalytic histidine and aspartate residues are indicated in red.

## Supplementary Table

**Supplementary Table 1.**
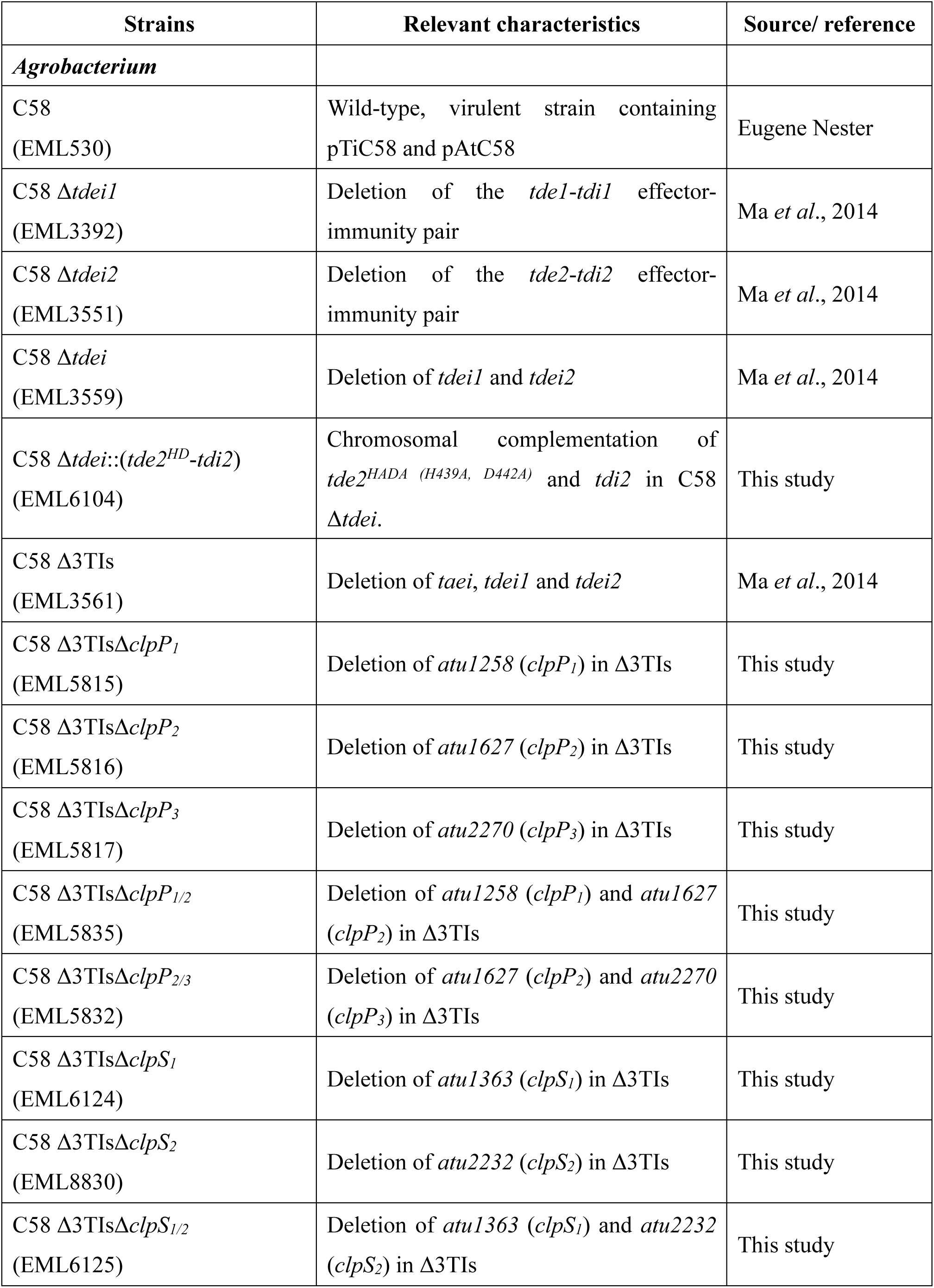
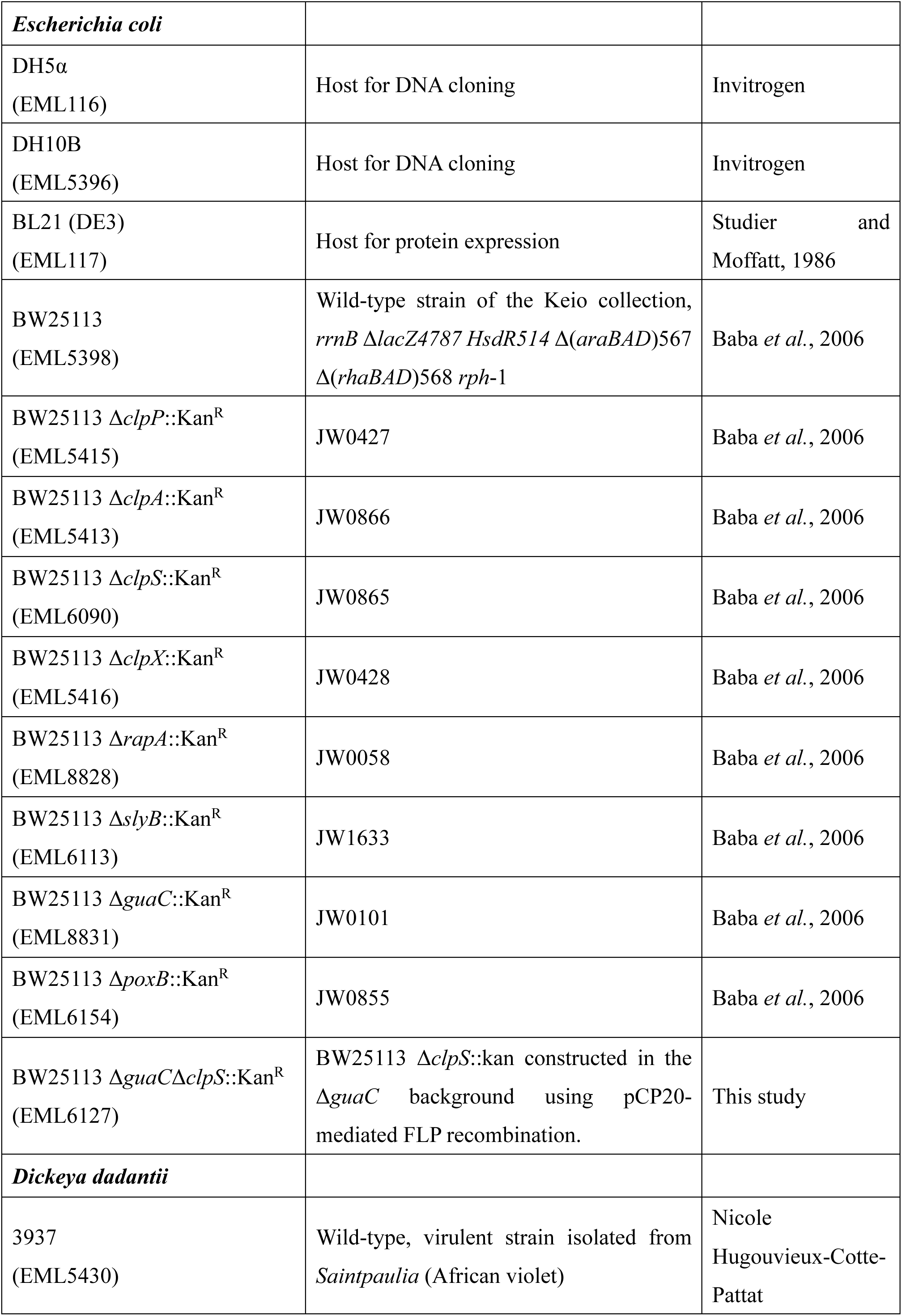
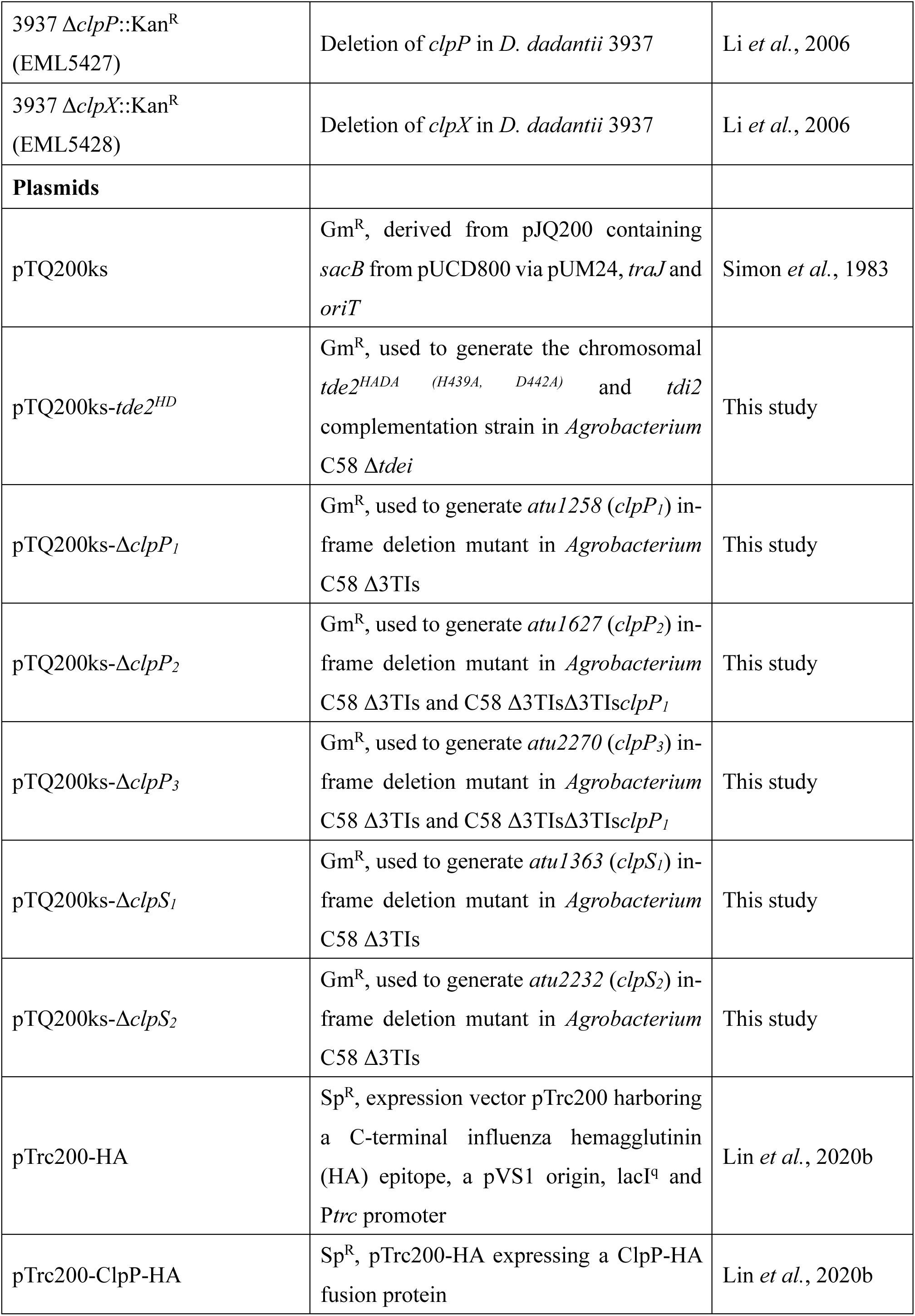
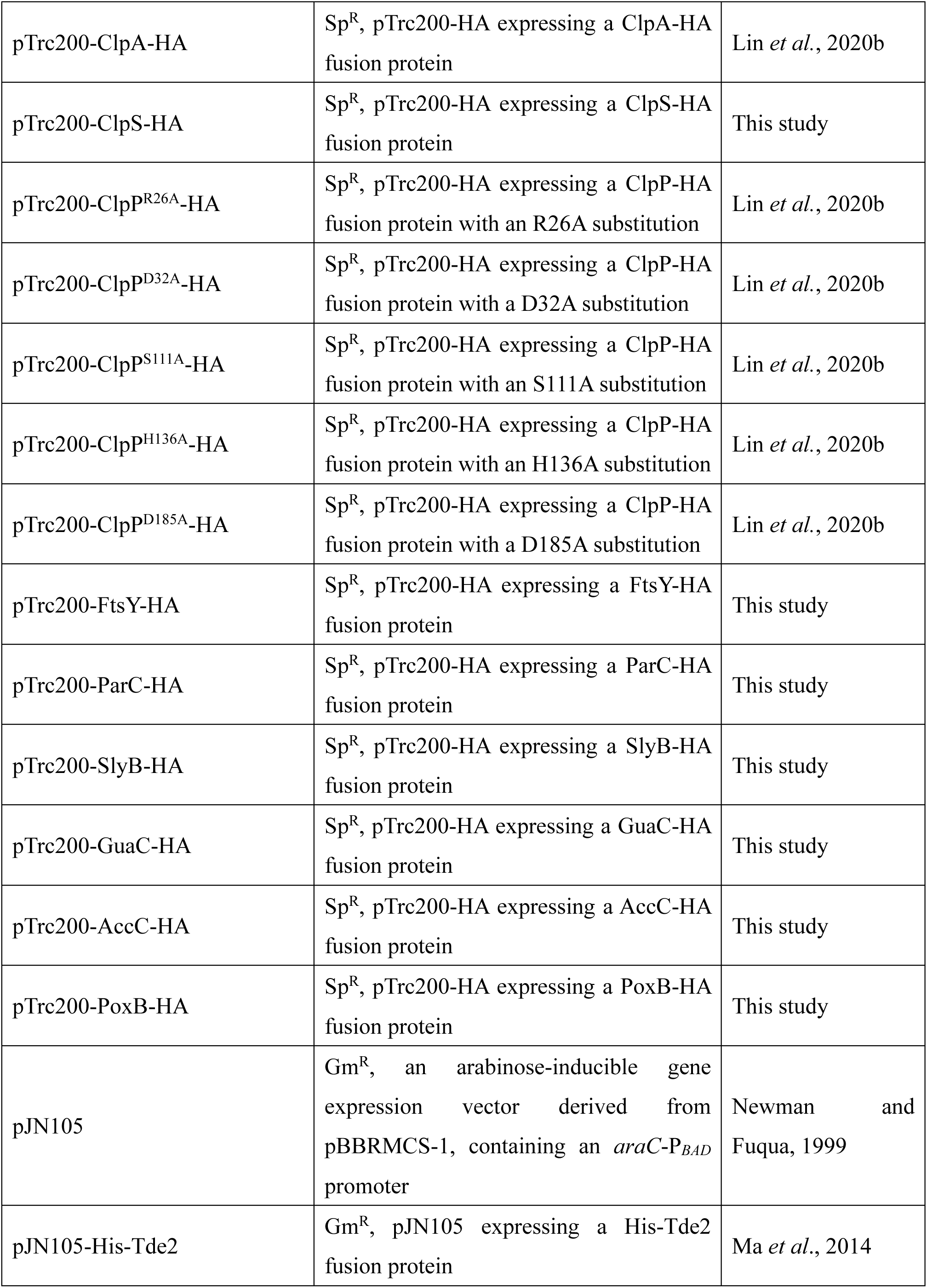
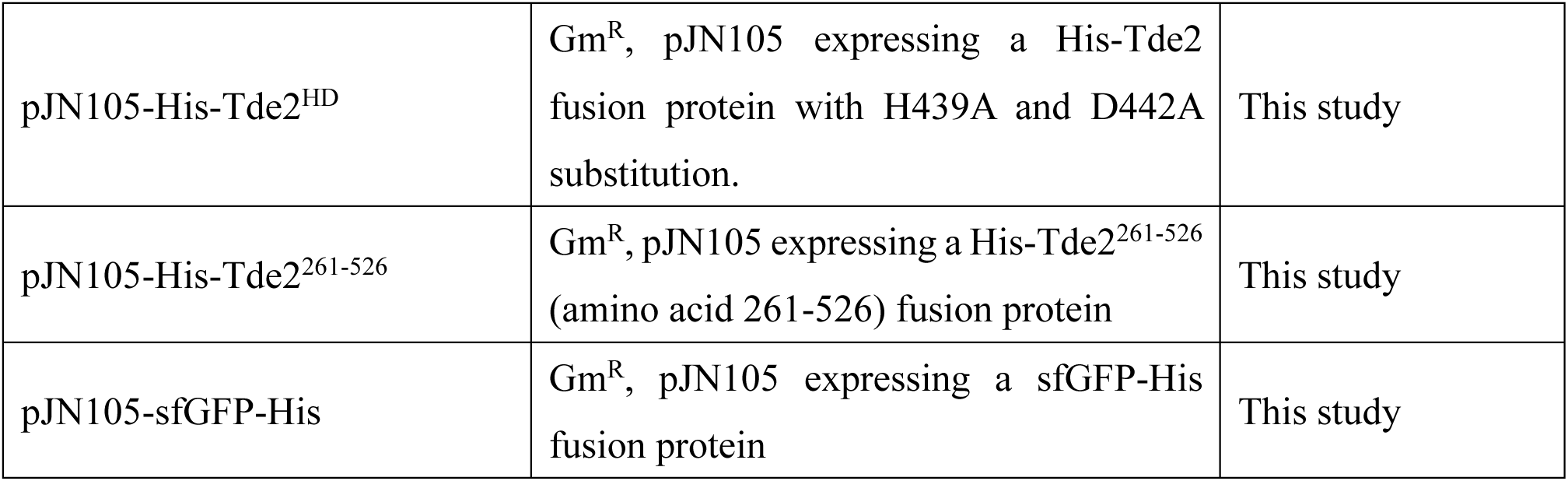
Bacterial strains and plasmids.

**Supplementary Table 2.**
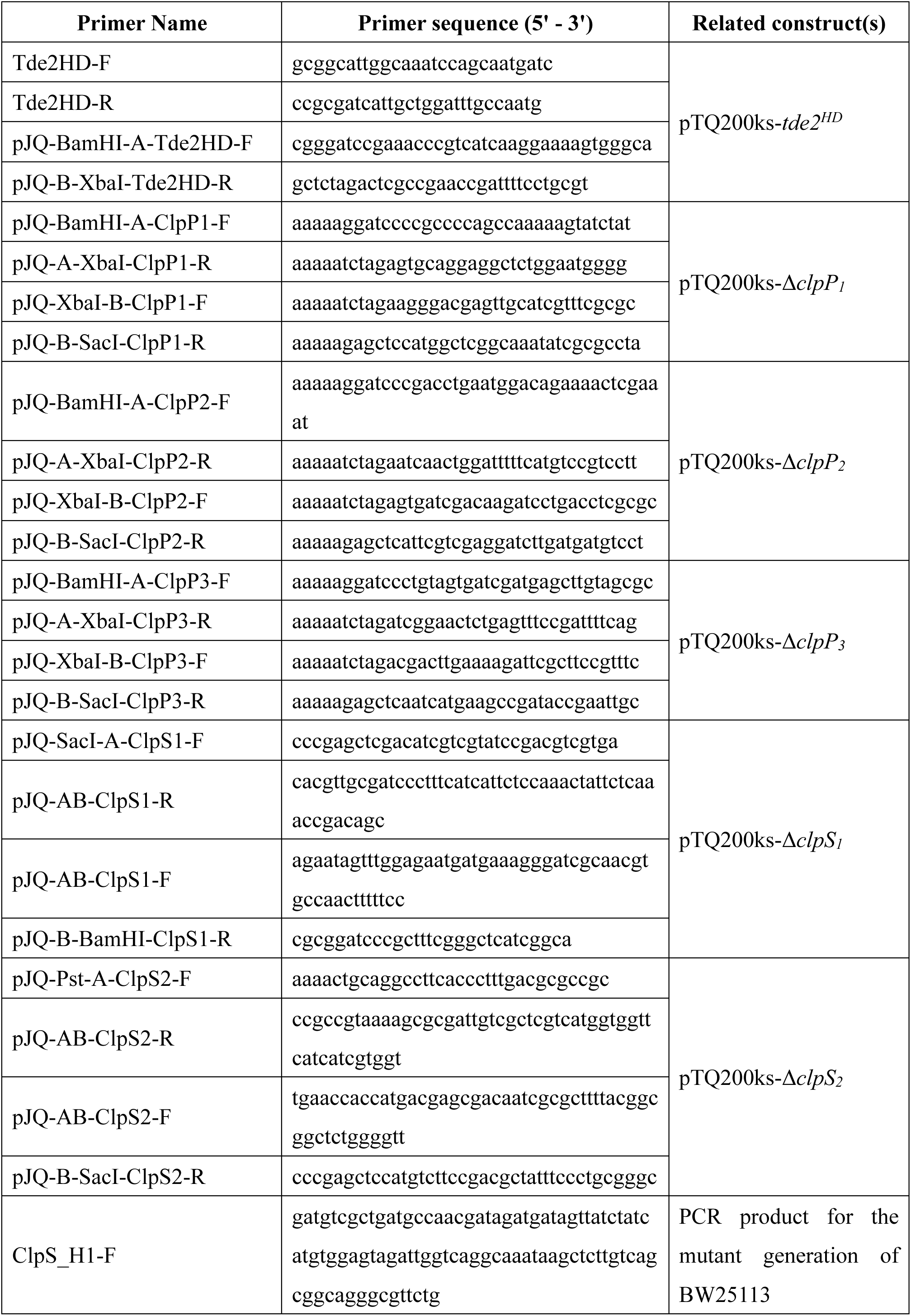
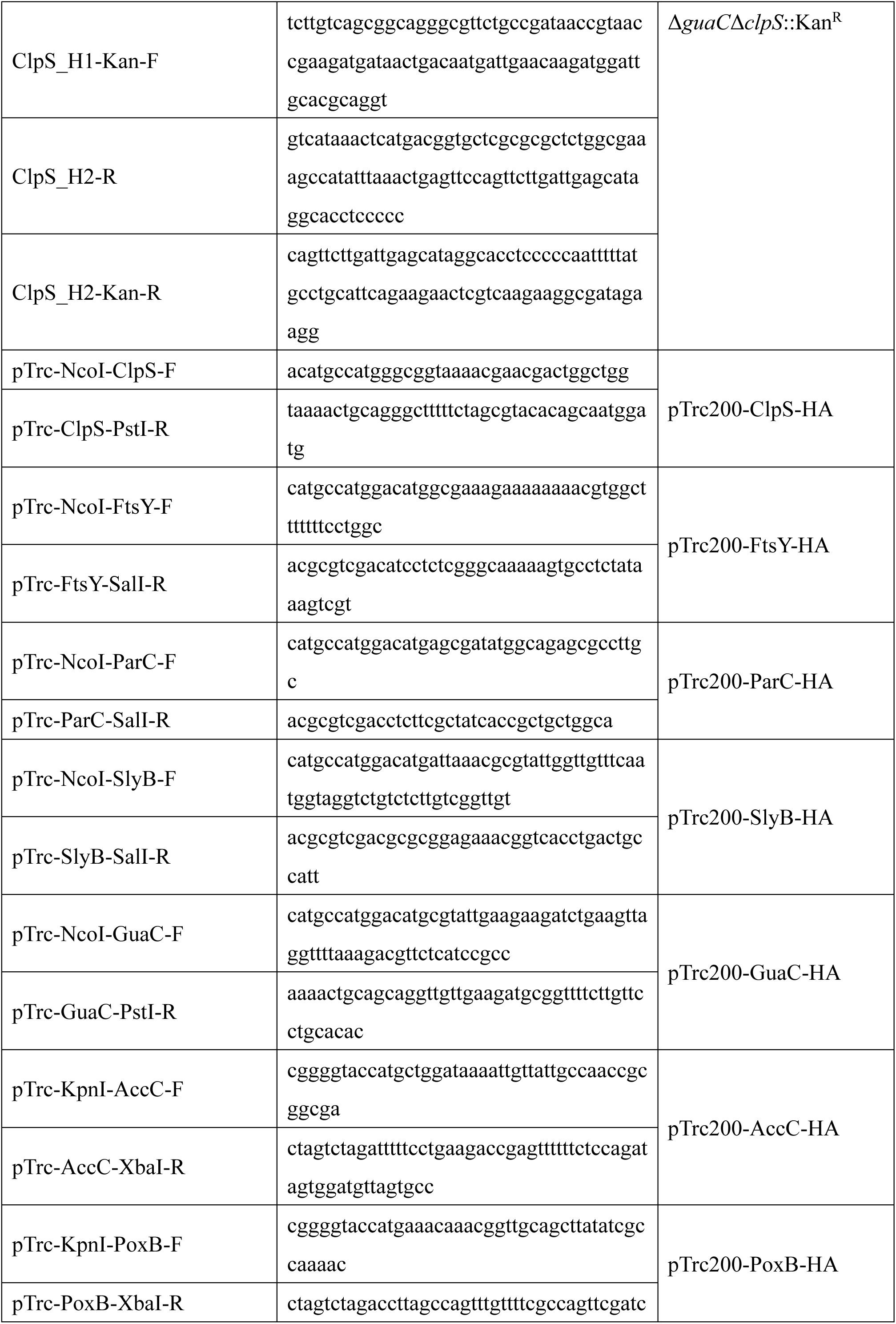
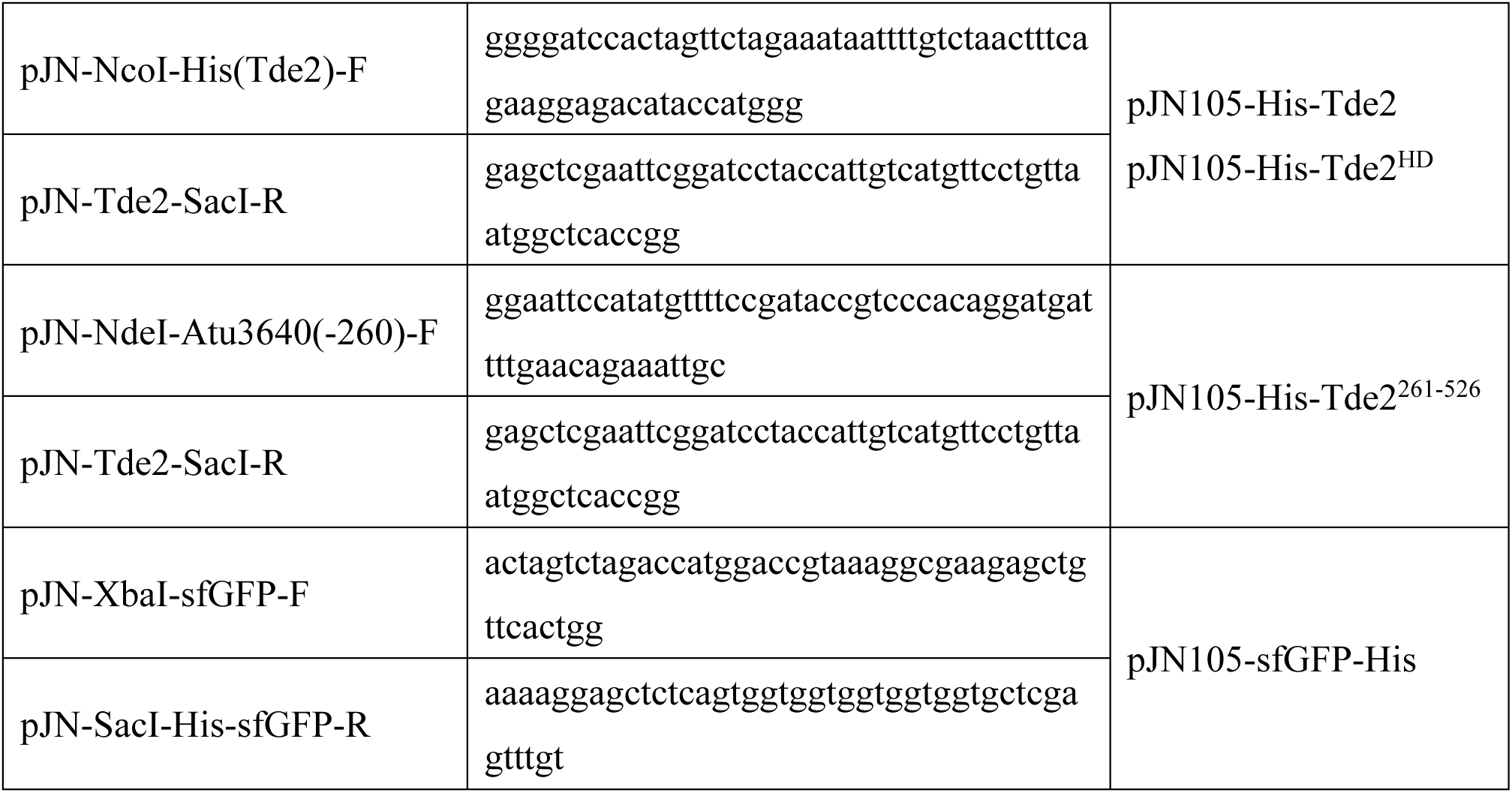
Primer list.

